# Machine learning prediction of enzyme optimum pH

**DOI:** 10.1101/2023.06.22.544776

**Authors:** Japheth E. Gado, Matthew Knotts, Ada Y. Shaw, Debora Marks, Nicholas P. Gauthier, Chris Sander, Gregg T. Beckham

## Abstract

The relationship between pH and enzyme catalytic activity, especially the optimal pH (pHopt) at which enzymes function, is critical for biotechnological applications. Hence, computational methods to predict pHopt will enhance enzyme discovery and design by facilitating accurate identification of enzymes that function optimally at specific pH levels, and by elucidating sequence-function relationships. In this study, we proposed and evaluated various machine-learning methods for predicting pHopt, conducting extensive hyperparameter optimization, and training over 11,000 model instances. Our results demonstrate that models utilizing language model embeddings markedly outperform other methods in predicting pHopt. We present EpHod, the best-performing model, to predict pHopt, making it publicly available to researchers. From sequence data, EpHod directly learns structural and biophysical features that relate to pHopt, including proximity of residues to the catalytic center and the accessibility of solvent molecules. Overall, EpHod presents a promising advancement in pHopt prediction and will potentially speed up the development of enzyme technologies.

## Introduction

Enzyme activity is significantly influenced by the pH of the reaction environment, often with a marked decline in activity beyond a specific pH range due to catalytic deactivation and structural instability.^1,2^ While most characterized enzymes have an optimal pH for activity (pHopt) close to the neutral value of 7.0, some enzymes function optimally at extremely acidic or alkaline conditions, with acidic or alkaline pHopt values as low as 1.0 or as high as 12.5, respectively.^3–5^ In industrial biochemical processes, enzymes are frequently utilized, or desired to be used, far from their pHopt, leading to a substantial reduction in activity. As a result, there is a growing interest in discovering and engineering enzymes with enhanced pH tolerance to overcome these limitations.

Exploring genomes from organisms or metagenomes from environments with extreme pH values, based on sequence homology with characterized enzymes, is a promising method for identifying natural enzymes with improved pH resistance.^5–7^ However, enzymes identified through this approach may lack sufficient catalytic activity.^8^ Alternatively, an enzyme with proven activity can be modified to adjust its pH-activity profile through methods such as immobilization,^9,10^ chemical modification,^11–13^ or protein engineering.^14,15^ Machine learning significantly benefits these approaches by enabling accelerated discovery and accurate prediction of enzymes with a desired pHopt, effectively learning complex relationships between enzyme sequence and pH. Recently, researchers have employed computational approaches to investigate protein pH relationships with biophysical methods,^16–18^ and to predict enzyme pHopt using traditional machine learning models with limited datasets (fewer than 500 proteins).^7,19–25^ Despite these efforts, the adoption of large-scale machine learning to capitalize on recent breakthroughs in the field remains limited, primarily due to the scarcity of sufficiently large enzyme pH datasets for training purposes.

In this work, we introduce two pH optima datasets (enzyme catalytic and organism environment optima) to advance learning protein sequence-pH relationships. We extensively train and evaluate various machine learning methods on these datasets, optimizing hyperparameters for each method. Our optimal method, EpHod, predicts enzyme pHopt directly from protein sequences utilizing embeddings from the ESM-1v protein language model, and achieves a root mean squared error (RMSE) of 1.25 pH units on the held-out test set. We show that EpHod effectively learned to focus on residues near the catalytic center that predominantly affect binding and catalytic activity, and on residues at the protein surface that interact with solvent molecules and ions and are crucial for maintaining structural stability in extreme pH environments. We further demonstrate that compared to existing biophysical methods, EpHod achieves more accurate prediction of catalytic pHopt, and maintains robust performance on enzyme sequences that differ significantly from those encountered during training.

## Results

### Optimal pH datasets for model training

From BRENDA,^26^ we compiled a dataset of 9,855 enzymes that are annotated with the pH values at which optimal enzyme activity was measured experimentally. This dataset, which we refer to as the pHopt dataset, comprises enzymes from organisms across the tree of life (Figure 1C, Supplementary Table 1) and includes the seven main enzyme activity classes described by enzyme commission (EC) numbers, with hydrolases (EC 3.x.x.x) being the most prevalent class (Figure 1D). We divided the pHopt dataset into training, validation, and testing sets, containing 7124, 760, and 1971 sequences respectively. Sequences in the validation set had less than 20% identity with training sequences, while the testing set had a similar distribution as the training set to enable comprehensive comparative analyses over the full sequence distribution. A subset of the testing set (999 sequences) was curated to have less than 20% identity with training sequences, and was utilized to evaluate the predictive performance on sequences significantly different from training sequences.

**Figure 1.**
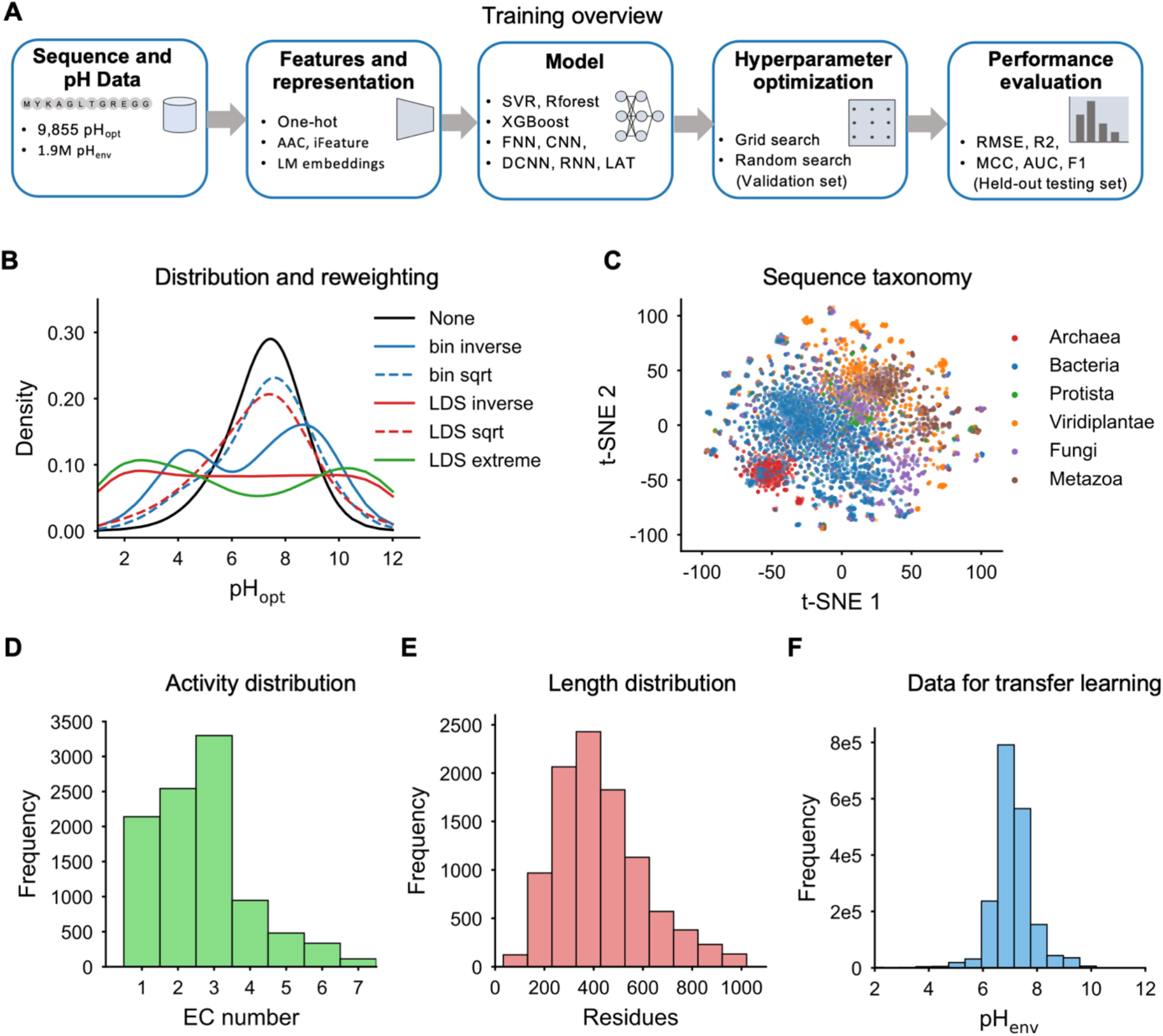
Overview of model training and the distribution of the training data. **(A)** Enzyme optimum pH (pHopt) and organism environment pH (pHenv) are used to train various machine learning methods with different sequence representations. For each method, up to 200 model instances are trained with varying hyperparameters, and the optimal model, selected by performance on the validation set, is assessed on a held-out testing set. **(B)** Kernel density estimate (KDE) of pHopt values for 9,855 enzymes before (None) and after applying reweighting strategies to address the imbalance of the label distribution (colored and dashed lines). **(C)** Taxonomical diversity (kingdom) of 9,855 pHopt enzymes visualized by a two-dimensional t-SNE reduction of averaged ESM-1v embeddings.^39^ **(D)** Distribution of enzyme commission (EC) numbers of 9,855 pHopt enzymes (1: oxidoreductases, 2: transferases, 3: hydrolases, 4: lyases, 5: isomerases, 6: ligases, 7: translocases). **(E)** Distribution of sequence length of 9,855 pHopt enzymes. **(F)** Distribution of pHenv values of 1.9 million secreted bacterial proteins used in transfer learning.

Given the limited size of the pHopt dataset, we hypothesized that pre-training models on a larger dataset of environment pH values would improve pHopt prediction by exploiting the observed association between environment pH and exoenzyme optimum pH (Supplementary Figure 1).^27,28^ Consequently, we prepared an additional dataset of 1.9 million secreted bacterial proteins from BacDive and NCBI databases,^29,30^ that are mapped to the environmental pH values for optimal growth of the source organisms. We selected only secreted proteins by predicting the presence of a signal peptide with SignalP 6.0,^31^ since microbes maintain intracellular pH close to neutrality through homeostatic mechanisms.^2,32–34^ We used this dataset, which we refer to as the pHenv dataset, to examine the potential advantages of transfer learning,^28,35^ by comparing models pre-trained on pHenv values and then fine-tuned on pHopt values, with models directly trained on pHopt values. To avoid data leakage in pre-training, we selected the pHenv training dataset (1.71 million) to share less than 20% identity with the pHopt testing set.

Previous studies have demonstrated that training with imbalanced datasets can result in overfitting of the models towards abundant regions leading to large errors on sparse regions of the target label values.^36,37^ Our analysis revealed that both pHopt and pHenv datasets exhibit highly imbalanced label distributions (Figures 1B and 1F). Nearly 75% of the pHopt values lie between pH 6 to 8, and 94% are between pH 5 and 9. Considering that accurate prediction of extreme acidic and alkaline enzymes would be especially desirable for biotechnological applications, we addressed the label distribution imbalance by employing several techniques during model training to reweight the loss function with values inversely proportional to the label density near each sample, thus directing the models to focus on sparse extreme regions. During hyperparameter optimization for each machine learning method, we selected the optimal reweighting technique from five approaches that derive sample weights by computing the inverse of the sizes of binned target values (bin techniques) or the inverse of a kernel-smoothed distribution of label values (label distribution smoothing, or LDS, techniques).^38^

### Models utilizing language model embeddings outperform alternative machine learning methods

To determine the most effective methods for predicting enzyme pHopt, we conducted an extensive evaluation of multiple machine learning models using various numerical representations of protein sequences. These representations included one-hot encodings, amino acid composition (AAC), an extensive set of engineered expert features from the iFeature package,^40^ and embeddings derived from protein language models (PLM). These PLMs, trained in a self-supervised manner, with the masked or next-token objective, are known to yield embeddings that richly capture protein properties.^41^ For this study, we selected a range of PLMs from the literature, including ESM-1b,^42^ ESM-1v,^39^ ProtT5,^43^ Tranception,^44^ ProGen2,^45^ and CARP (Supplementary Table 2).^46^

For the machine learning methods, we employed traditional machine learning models including ridge regression, k-nearest neighbor regression (KNR), support vector regression (SVR), random forest regression (RForest), and XGBoost. Additionally, we explored neural network architectures including fully-connected neural networks (FNN), convolutional neural networks (CNN), dilated convolutional neural networks (DCNN), and recurrent neural networks (RNN). Specifically for per-residue PLM embeddings, we trained a light attention model (LAT),^47^ and proposed several variations to LAT, including residual light attention (RLAT), perceptive attention (PAT), and dilated convolution light attention (DCAT). We trained these attention-based neural networks only on ESM-1v per-residue embeddings because of the significant computational costs associated with deep neural networks. We also chose ESM-1v among the PLMs because its averaged embeddings yielded the best performance on the pHopt validation set (Supplementary Figure 2).

Across the different combinations of numerical representations and model types, we trained up to 200 model instances per combination, each with a unique set of hyperparameters that were sampled via grid or random search to explore the hyperparameter space.^48^ This extensive training resulted in training 11,550 models in total (Supplementary Tables 3 to 6). We selected the optimal model hyperparameters based on performance on the validation set, and evaluated the optimal models on the held-out pHopt testing set. The performance of the optimal models on the held-out testing set is shown in Figure 2 and Supplementary Figures 3 to 6. Details of the machine learning approaches are provided in the Methods.

**Figure 2.**
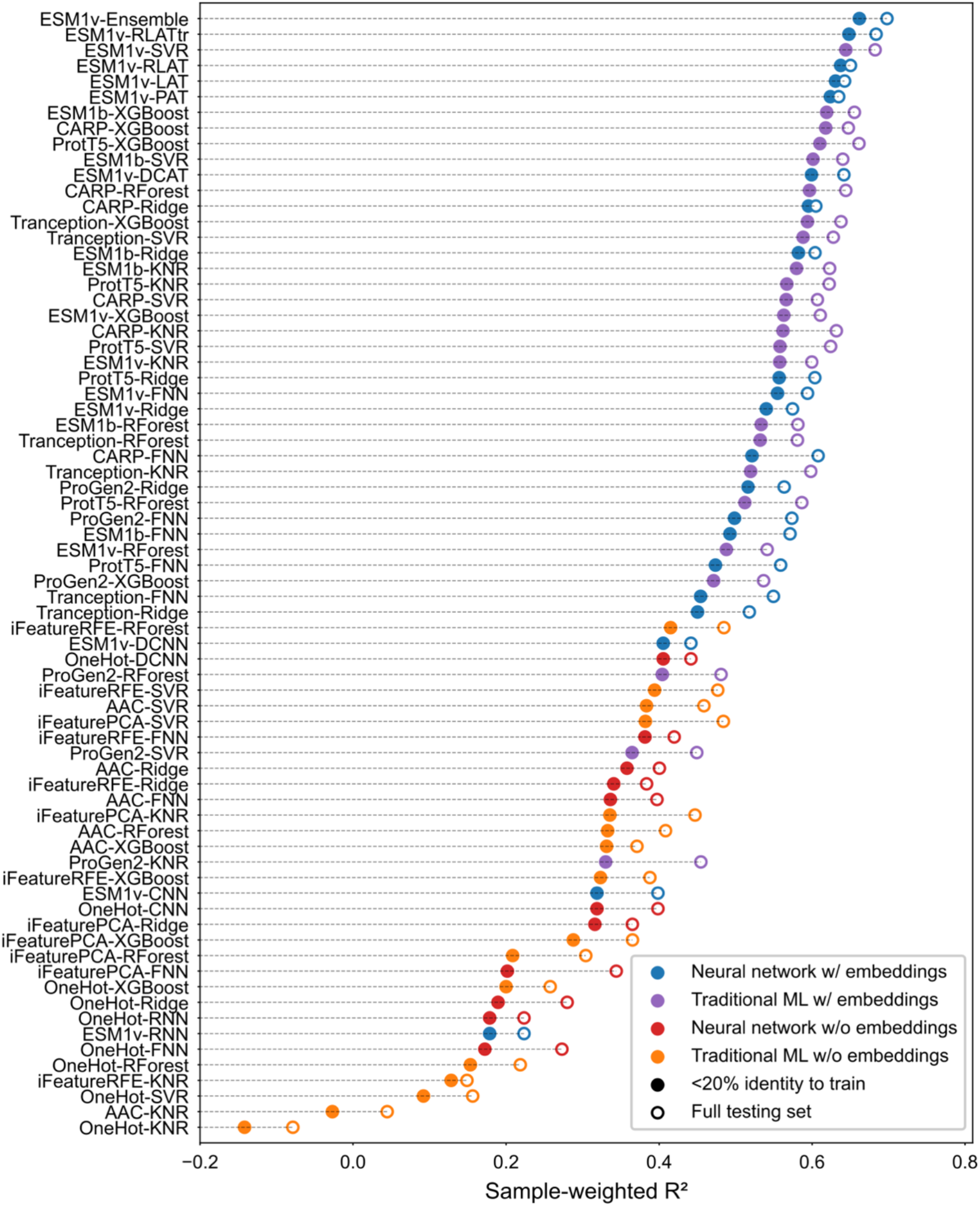
Performance of optimal models from each method on the complete held-out pHopt testing set (n=1971) and on a subset of the testing set with less than 20% sequence identity to the training set (n=999). Performance is quantified using the coefficient of determination (R^2^) of predicted pHopt values, adjusted by *bin-inverse* sample weights to adequately capture predictions in the sparse acidic and alkaline regions of the pHopt data. The y-axis labels indicate the sequence representation and model type used, respectively. For each combination of representation and model, up to 200 model instances were trained with varying hyperparameters. The optimal model was chosen based on the performance on a separate validation set. See Supplementary Figures 3 to 6 for RMSE, AUC, MCC, and F1 score.

The evaluation results demonstrate that methods utilizing PLM embeddings markedly outperformed those using other representations, achieving substantially higher R^2^ values on the held-out pHopt testing set. Notably, the best-performing method without language model embeddings combined random forest with feature selection (recursive feature elimination) of the iFeature representations (iFeatureRFE-RForest). This method achieved an R^2^ of 0.415 on testing set sequences with less than 20% identity to the training set. In contrast, an SVR model trained with averaged ESM-1v embeddings achieved an R^2^ of 0.644, representing a 55% improvement, and underscoring the superiority of language model embeddings. Furthermore, it is noteworthy that the iFeatureRFE-RForest approach also outperformed all neural network models that did not utilize PLM embeddings. This finding is consistent with a previous study that reported no performance advantage of deep learning over expert feature engineering in the prediction of enzyme optimal temperatures.^49^

### Transfer learning provides modest improvement in performance

The best-performing method utilizing PLM embeddings and neural networks employed a residual light attention model with ESM-1v per-residue embeddings (ESM1v-RLAT). To investigate potential performance improvements, we implemented transfer learning with the RLAT architecture and ESM-1v (Figure 3A). Specifically, we first pre-trained the model on the pHenv dataset followed by fine-tuning on the pHopt dataset (Figures 3C and 3D). Due to computational limitations and the risk of overriding valuable information acquired during self-supervised training,^50^ we elected to keep the ESM-1v model frozen during both pre-training and fine-tuning stages.

**Figure 3.**
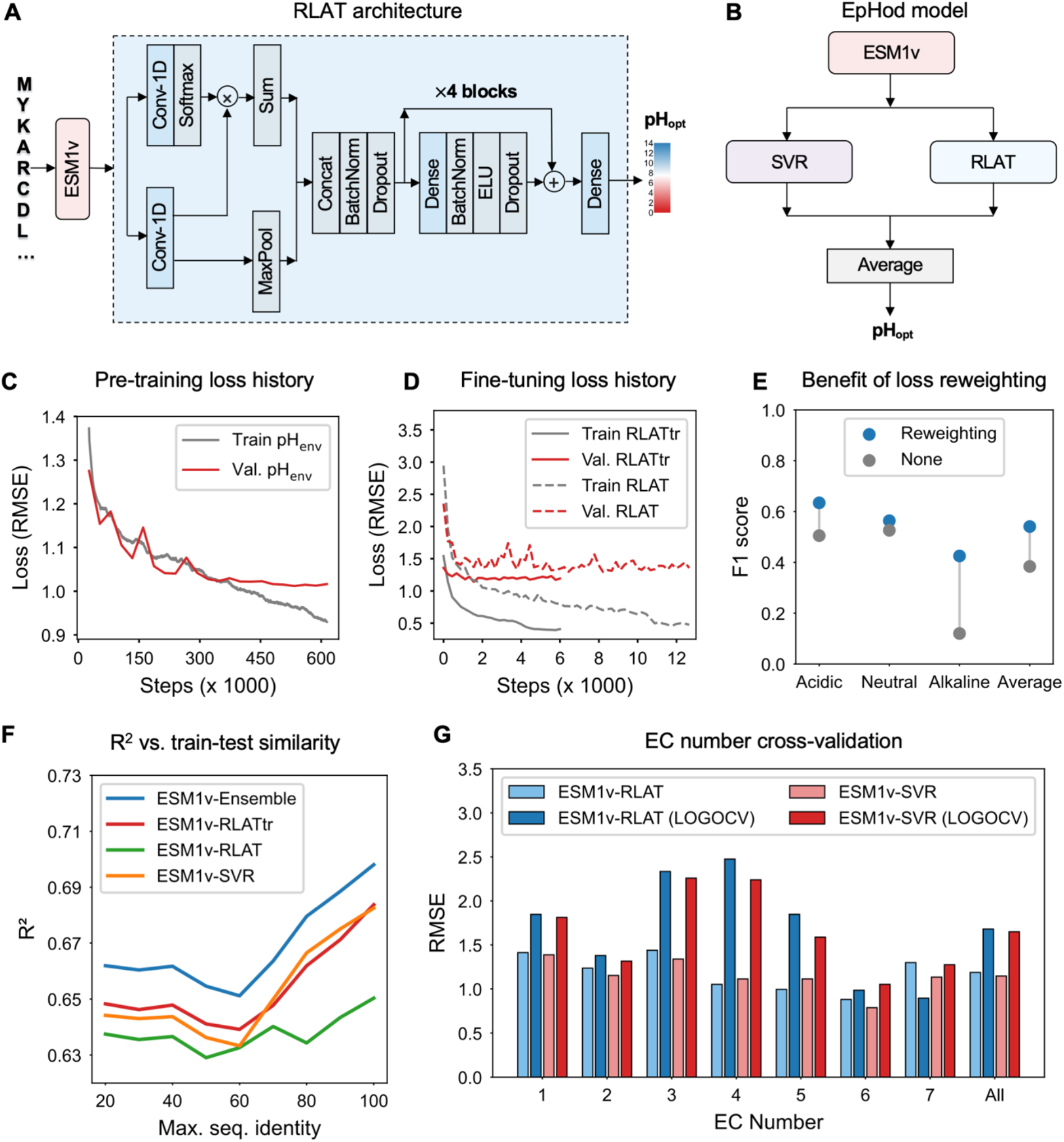
EpHod training and evaluation of performance. **(A)** Model architecture of the residual light attention (RLAT) model with four residual blocks of dense layers. The RLAT model is trained on top of frozen per-residue ESM-1v embeddings to predict enzyme pHopt. (**B**) EpHod is an ensemble of RLAT and support vector regression trained with per-residue and averaged ESM-1v embeddings, respectively. (**C**) RMSE loss of the RLATtr model during pre-training on the pHenv dataset. The training loss is shown as a moving average over one epoch or 26,688 steps, and validation loss is shown per epoch. (**D**) Loss during fine-tuning on the pHopt dataset after pre-training on the pHenv dataset (RLATtr). Dashed curves show the loss of the RLAT model trained directly on the pHopt dataset without pre-training (**E**) Effect of loss reweighting on the performance of the RLAT model. (**F**) Performance of EpHod (ensemble) and base models with respect to sequence similarity between testing and training sequences. (**G**) Sample-weighted RMSE evaluated with leave-one-group-out cross-validation (LOGOCV) with respect to enzyme commission numbers.

The transfer learning approach (ESM1v-RLATtr) demonstrated a modest performance improvement over the ESM1v-RLAT model trained exclusively on pHopt data, with a 5% enhancement in R2 score and RMSE (Figures 2 and 3F). However, the performance of the ESM1v-RLATtr model was only comparable to that of an SVR model trained with averaged ESM1v-embeddings (ESM1v-SVR), with R2 scores of 0.648 and 0.644, respectively, on testing set sequences with less than 20% identity to the training set. We also assessed an ensemble of the ESM1v-SVR and ESM1v-RLATtr models, which resulted in further performance gains, with an R2 score of 0.662. We refer to this ensemble approach (Figure 3B), which achieved the best overall performance, as EpHod (enzyme pH optimum prediction with deep learning).

We further compared EpHod with a recently published model, OphPred, by training OphPred on our pHopt data.^51^ While the authors report improved mean absolute error compared to EpHod (0.6 vs. 0.7), we note that this is largely dominated by the abundant neutral pHopt values in the data. We find that based on sample-weighted metrics that prevent overestimating performance on neutral pH values, but equally emphasize sparse acidic and alkaline regions, EpHod outperforms OphPred (Supplementary Figure 7).

### EpHod generalizes across diverse enzyme sequences

To comprehensively evaluate EpHod’s performance, we investigated the impact of the training approach on predictive accuracy and the model’s ability to generalize across diverse sequences relative to the training set. Given that deep learning models have been reported to perform worse than random predictions on proteins with less than 50% identity to the training sequences,^49,52^ we evaluated the performance of EpHod in relation to the similarity between testing and training sequences as a way to assess whether the predictive performance was majorly a consequence of high sequence similarity. As expected, EpHod and similar models exhibited a slight reduction in performance as sequence identity decreased between the training and testing sets (Figures 2 and 3F, Supplementary Figure 8). However, this decline was minimal, with an increase in RMSE of less than 0.12 pH units, indicating that EpHod maintains robust performance even on sequences with low identity to those in the pHopt training set.

Moreover, we evaluated the influence of the pHopt data distribution on the model performance and examined the benefits of reweighting the loss function during training. To this end, we compared the performance of the ESM1v-RLAT model, trained with sample weights inversely proportional to the size of binned label values (*bin-inverse*), to a similar model trained without reweighting. We used the RLAT model, rather than RLATtr, for this analysis to isolate the effects of the pHopt data distribution without the confounding impact of pre-training on the pHenv dataset. As anticipated, training without loss reweighting led to substantially poorer performance in the sparse acidic (pH<5) and alkaline (pH>9) range, as measured by the F1 score (Figure 3E). In contrast, reweighting the loss function resulted in 26% and 250% improvement in F1 score for acidic and alkaline enzymes, respectively, highlighting the effectiveness of this approach in addressing the scarcity of data in these regions. It is noteworthy that during hyperparameter optimization, we observed that the choice of reweighting technique (Figure 1B) had a negligible effect on performance, and in most cases, simply reweighting based on the inverse of bin sizes (*bin-inverse*) was sufficient for optimal results.

Furthermore, we evaluated the variability in predictive performance with respect to the seven main enzyme commission numbers, assessing EpHod’s ability to generalize to enzyme activities not represented during training. We compared the errors of ESM1v-RLAT and ESM1v-SVR models for enzymes from each EC number against errors obtained from leave-one-group-out cross-validation, where similar models were trained on modified training sets excluding sequences from each EC number. The RMSE values ranged from 0.78 to 1.44 pH units when all EC numbers were included in training, but excluding sequences from specific EC numbers led to a moderate increase in error, up to 2.48 pH units. Notably, the SVR model demonstrated superior generalization to EC numbers absent from the training set compared to the RLAT neural network (Figure 3G). Altogether, despite the variability in performance, these findings indicate that EpHod effectively learns features relevant to pHopt across diverse enzyme activities, enabling it to generalize well to underrepresented or unseen activity classes.

### EpHod captures relevant structural features related to enzyme pH optima

The relationship between protein structural features and enzyme pH optima is well-documented in literature.^2^ The local distribution of amino acids in the protein structure impacts stability and crucial catalytic interactions under various pH conditions.^53^ Although a universal mechanism remains elusive, the adaptation of acidic and alkaline proteins to extreme pH environments is often attributed to an abundance of negatively charged (Asp, Glu) and positively charged amino acids (Arg, Lys) on the protein surface.^2,54,55^ Multiple protein engineering studies have demonstrated that altering the composition of charged residues to increase the net negative or positive charge, particularly on the protein surface, can shift the pHopt towards more acidic,^56–58^ or more alkaline values,^59–61^ respectively. Moreover, residues in closer proximity to catalytic sites exert greater influence on the atomic environment and protonation states (pKa) of catalytic residues, thereby dominantly affecting the pH optima.^15,62^ Consequently, mutating residues near the catalytic residues to alter the pKa of catalytic residues and maintain their catalytic protonation states at extreme pH levels is recognized as a viable method for shifting the pHopt.^14^

Based on these relationships, we next examined whether EpHod acquired these structural features during gradient-based training on the pHenv and pHopt datasets. While several interpretability methods are available,^63^ in this work, we assessed the attention weights per residue in the RLATtr top model (output of the softmax layer) to identify the residues EpHod emphasizes for inference (Figure 3A, details in Methods). We observed that certain residues in the proteins exhibited considerably greater attention weights, suggesting that these residues play a prominent role in pHopt prediction (**Figure 4B**). However, it is important to note that the contextual information inherent in the ESM-1v per-residue embeddings from the self-attention mechanism may influence the interpretation of these attention weights, since some information from surrounding residues are captured in each residue’s embedding.

**Figure 4.**
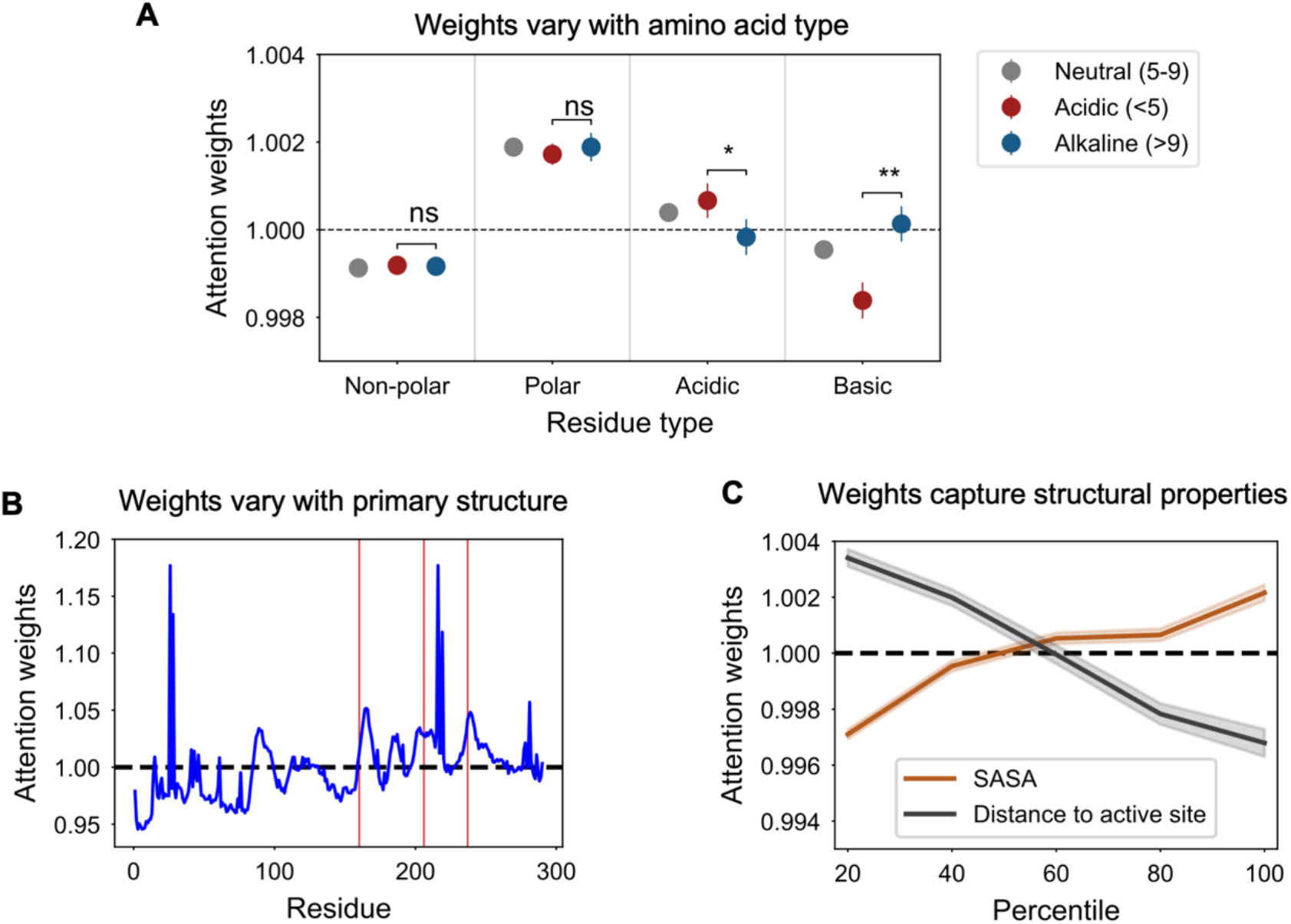
Analysis of attention weights in EpHod’s RLAT model revealing physicochemical and structural features associated with pHopt using the full dataset (n=9,855). The dashed line (1.0) indicates mean attention weights per protein after normalization. **(A)** Average attention weights of residues classified as non-polar (Asp, Val, Leu, Ile, Pro, Phe, Trp, Met, Gly), polar (Ser, Thr, Cys, Tyr, Asn, Gln), acidic (Asp, Glu), and basic (Arg, Lys, His). Error bars indicate 95% confidence intervals of the mean. A two-sided Welch’s t-test comparing average weights in acidic and alkaline enzymes reveals statistically significant difference in acidic and basic residues with *p*-values of 4.1E-3 (^*^) and 3.4E-9 (^**^), respectively, but non-significant difference in non-polar and polar residues, with p-values of 0.87 and 0.43, respectively. **(B)** Average attention weights of residues in primary structure of an example enzyme, *Ideonella sakaiensis* PETase (UniProt ID: A0A0K8P6T7).^64^ Catalytic residues (Ser160, Asp206, and His237) are indicated by red lines. Certain residues exhibit weights substantially greater than average in the model, contributing majorly to the predicted pHopt. **(C)** Averaged attention weights of residues compared with solvent accessible surface area (SASA) and distance from catalytic residues. To enable comparison across different proteins, SASA and distance values were binned into quintiles, and averaged values in each quintile are shown. The colored regions around the lines represent 95% confidence interval of the mean.

Intriguingly, we found that on average, polar residues were assigned higher attention weights than non-polar residues, with the weights being statistically indistinguishable for acidic and alkaline enzymes. Nonetheless, EpHod learned to selectively concentrate on negatively charged (acidic) residues in acidic enzymes and positively charged (basic) residues in alkaline enzymes, assigning significantly higher weights to acidic residues in acidic enzymes than alkaline enzymes, and higher weights to basic residues in alkaline enzymes than acidic enzymes (Figure 4A). Moreover, comparing the difference between acidic and alkaline enzymes, the rank order of individual amino acids by attention weights is considerably different from the rank order of the amino acid composition (AAC), indicating that EpHod learns the relative importance of amino acids beyond the distribution of AAC in the data (Supplementary Figure 9-10). We also evaluated the average weights in relation to solvent accessibility and proximity to catalytic residues. We found that, on average, EpHod allocated progressively higher weights to more exposed residues and to residues nearer to the catalytic center (Figure 4C), in accordance with experimental observations. This trend is also discernible when the weights are projected on structures of selected enzymes (Figure 5).

**Figure 5.**
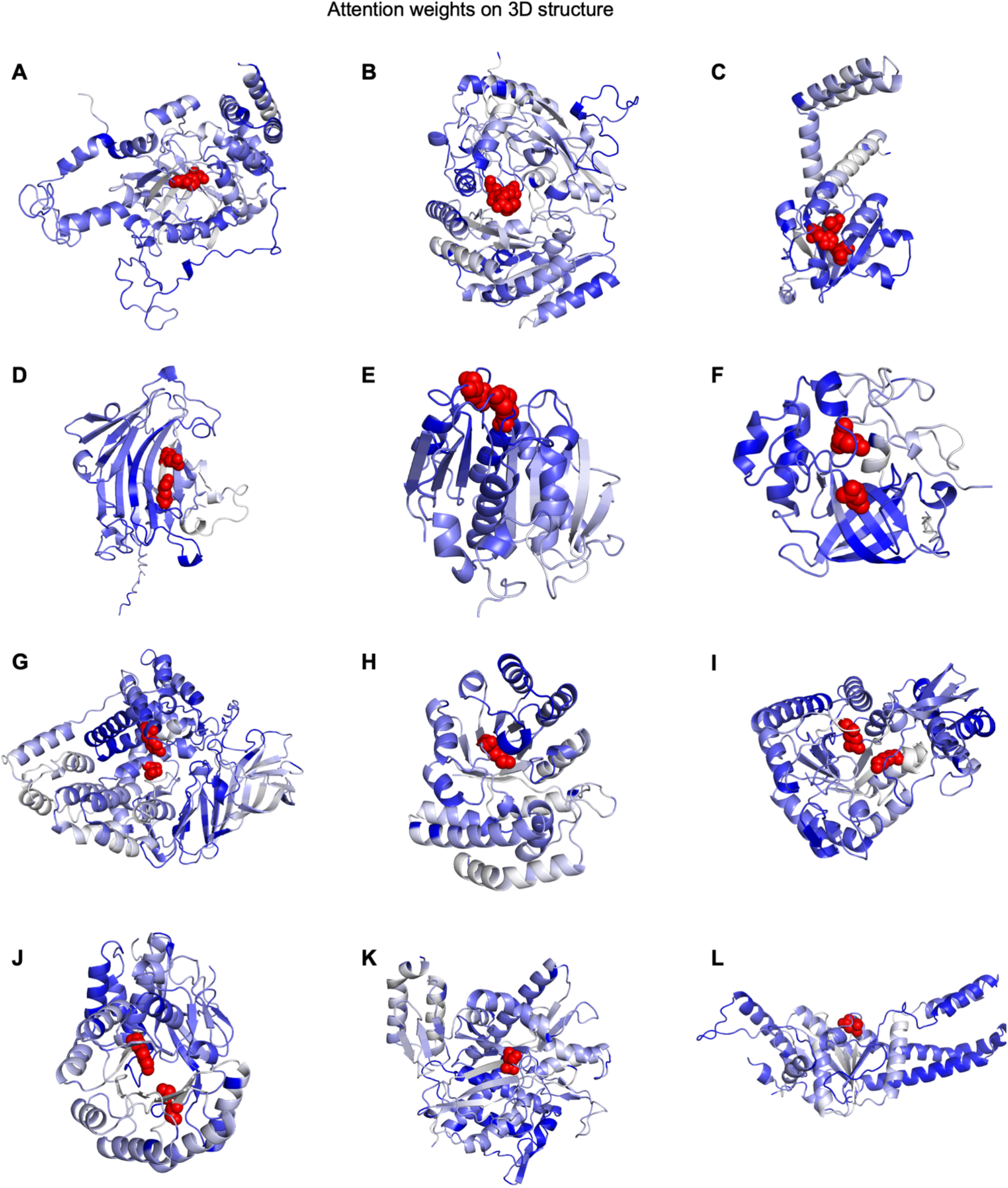
Visualization of per-residue attention weights of EpHod’s RLAT model on selected enzyme protein structures. Darker blue regions signify residues with higher weights, highlighting areas in the protein that EpHod emphasizes for pHopt predictions. Atoms of catalytic residues are shown in red. Weights were clipped to 0.5 and 99.5 percentiles to minimize the effects of outliers on the plot. Data in parenthesis indicate UniProt accession ID (UID), enzyme commission number, and the reported experiment pHopt in BRENDA, respectively. (**A)** *H. discus discus* catalase (UID: A5XB38, EC: 1.11.1.6, pHopt: 10.5), **(B)** *P. simplicissimum* vanillyl-alcohol oxidase (UID: P56216, EC: 1.1.3.38, pHopt: 7.0), **(C)** *H. sapiens* lecithin retinol acyltransferase (UID: O95237, EC: 2.3.1.135, pHopt: 9.1), **(D)** *D. kaki* xyloglucan endotransglucosylase (UID: G5DAC7, EC: 2.4.1.207, pHopt: 5.2), **(E)** *I. sakaiensis* poly(ethylene terephthalate) hydrolase (UID: A0A0K8P6T7, EC: 3.1.1.101, pHopt: 9.0), **(F)** *C. antarticus* endoglucanase (UID: D3GDK4, EC: 3.2.1.4, pHopt: 3.5), **(G)** *P. telluritireducens* aminopeptidase (UID: A0A3S6IVW5, EC 3.4.11.1, pHopt: 7.2), (**H)** *P. mume* aldehyde lyase (UID: Q4JC35, EC: 4.1.2.55, pHopt: 6.3), **(I)** *B. abortus* phosphopyruvate hydratase (UID: Q2YPV0, EC: 4.2.1.11, pHopt: 8.5), **(J)** *B. subtilis* L-Ala-D/L-Glu epimerase (UID: O34508, EC: 5.1.1.20, pHopt: 8.5), **(K)** *C. glutamicum* pup-protein ligase (UID: Q8NQE1, EC: 6.3.1.19, pHopt: 7.5) **(L)** *C. elegans* ATPase (UID: P30632, EC:6.3.1.19, pHopt: 7.4).

Furthermore, we compared EpHod with structural and biophysical methods, examining the ability of these methods to accurately differentiate between acidic and alkaline enzymes. We evaluated the area under the receiver operating characteristic curve (AUC) and the correlation between predicted scores and experimentally determined pHopt values. We considered several alternative methods, including the relative amounts of positively charged to negatively charged residues on the protein surface (i.e., Arg + Lys – Asp – Glu), the isoelectric point (pI), and an estimation of the pH for optimum stability (ΔΔG pHopt) using the PROPKA3 package.^65^ Our results demonstrate that EpHod achieved better performance compared with these other approaches in the ability to predict catalytic pHopt on the held-out testing dataset (Figure 6). We note that although PROPKA3 computes the pH at which the protein folding free energy is minimum, the optimum pH for activity and stability are generally correlated,^66^ and the optimum pH for catalytic activity has greater biochemical significance, since an enzyme needs to be sufficiently stable to carry out its function, but catalysis may be abolished regardless of stability. Our results are also in agreement with previous studies that found little or no correlation between the isoelectric point and optimum pH.^8,66,67^

**Figure 6.**
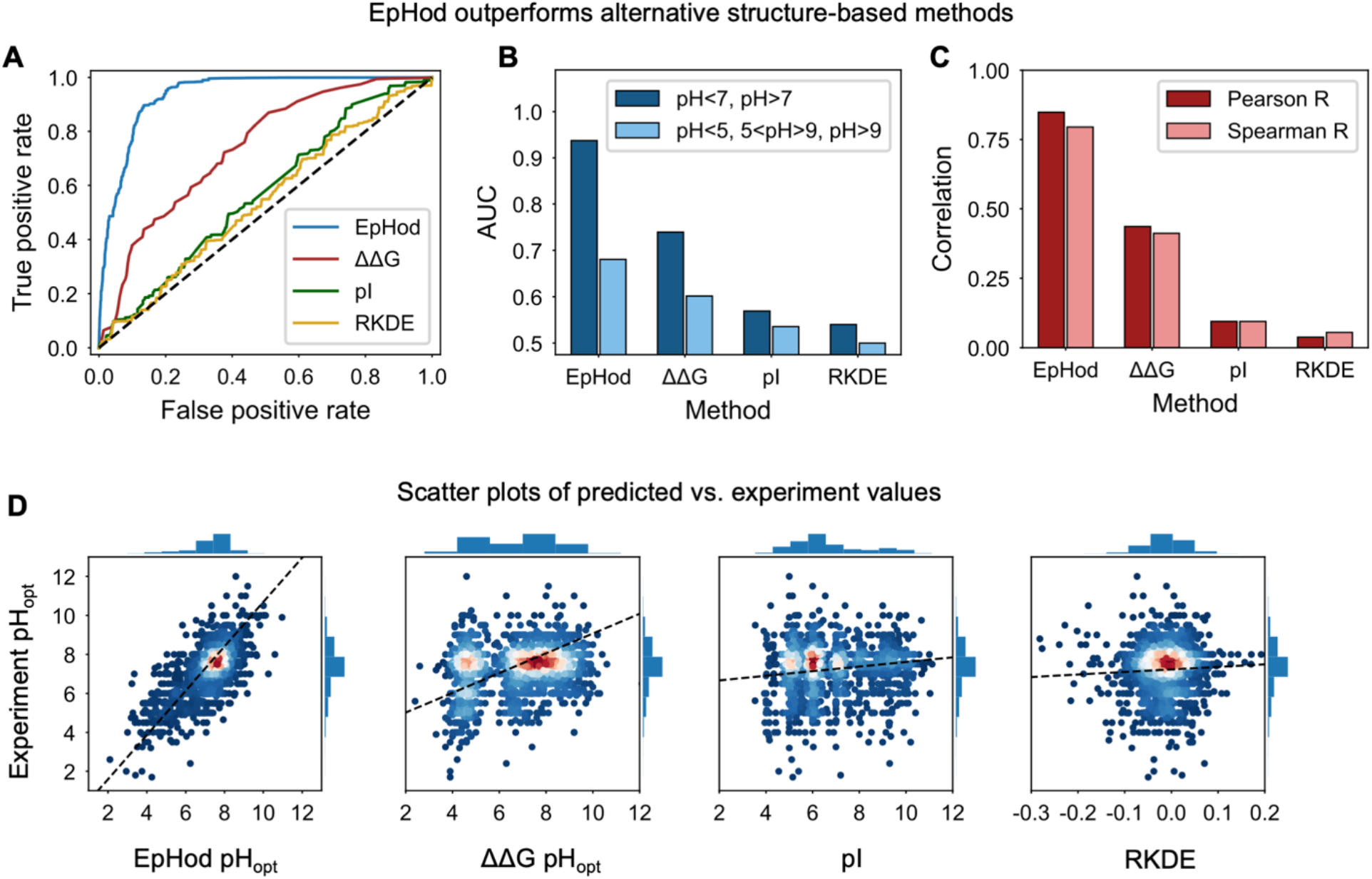
Comparison of the predictive performance of EpHod with alternative structural and biophysical methods on the full held-out testing set (n=1,971). In predicting pHopt values of the held-out testing dataset, EpHod outperforms calculations of the optimum pH for stability with PROPKA (ΔΔG pHopt),^65^ the isoelectric point (pI), and the relative amounts of positively charged amino acids (Arg and Lys) to negatively charged amino acids (Asp and Glu) on the protein surface (RKDE). **(A)** Receiver operating characteristic (ROC) curve showing true positive rate (TPR) against false positive rate (FPR) for a binary prediction of pHopt (pH<7, pH>7) at varying thresholds. The dashed line corresponds to random prediction and the area under each curve (AUC) is a measure of the predictive ability. **(B)** Area under the ROC curve for binary pHopt prediction, shown in **A**, and for a ternary prediction computed as the average of one-vs-rest AUC values. **(C)** Spearman and Pearson correlation coefficient between predictive score and experimental pHopt. **(D)** Scatter plots comparing predictive scores for each method and experimental pHopt. The color gradient of the points, ranging from blue to red, signifies low-to-high density. The dashed line represents a linear regression line fitted to a balanced dataset with respect to experiment pHopt distribution.

## Discussion

In this study, we present a machine learning model, EpHod, to predict enzyme pHopt values. Our extensive evaluations and comparisons of various machine learning methods clearly demonstrates the superiority of protein language model (PLM) embeddings for numerical representation of proteins, reinforcing the growing consensus that PLM embeddings drive state-of-the-art performance across various protein property prediction tasks.^31,47,68,69^. However, we observed considerable variability in the performance across different PLM embeddings, even after extensive hyperparameter optimization (Figure 2). This finding highlights the importance of carefully selecting the most suitable PLM embeddings for specific tasks when training predictive models. Moreover, embeddings from a CNN-based model, CARP, performed comparably to those from transformer based models, supporting the assertion from Yang et al. that CNNs can be competitive with transformer models when trained on the a similar scale of data and model size.^46^

Our results also indicate that when predicting properties of natural proteins with relatively smaller datasets (<10,000), traditional machine learning models, such as SVR and XGBoost, trained on averaged PLM embeddings, can perform on par or even outperform deep neural networks trained on full per-residue embeddings,^70^ although this finding may not extend to predicting effects of mutations on proteins.^71,72^ Even after pre-training on the larger pHenv dataset, we found that neural networks did not significantly outperform the traditional SVR approach. We surmise that recent advances in parameter-efficient fine-tuning,^73^ as well as multi-task learning,^74^ could potentially provide greater performance improvements compared to the classical fine-tuning we employed.

Moreover, our findings highlight the importance of considering the practical applications in which models will be utilized to ensure optimal training. Anticipating that the primary use of EpHod would be in identifying enzymes with high acid or alkaline tolerance, we adjusted the training process through a sample-weighted loss function and evaluated models using sample-weighted metrics. This approach mitigated biases toward prevalent neutral pH values and ensured the development of a model that excels in predicting extreme pH values. Beyond the pHopt distribution, EpHod also performs consistently well across enzymes of varying residue length, taxonomy, sequence similarity, and activity class, yielding RMSE values of less than 2.0 pH units for all analyzed categories (Supplementary Figure 11). Unlike many supervised models that exhibit substantial variability over sub-classes of the datasets,^36,52,75,76^ our results, and those of several recent studies,^31,68,70^ indicate that models employing PLMs more effectively address issues of data scarcity and imbalance, leveraging the information garnered from self-supervised pretraining.^77^

Furthermore, we anticipate that EpHod will offer multiple applications in enzyme discovery and design. First, EpHod could be used for high-throughput predictions of pHopt for enzyme homologs from extensive sequence databases,^78,79^ enabling identification of enzymes with enhanced acid or alkaline tolerance. Second, residues yielding large attention weights, or point mutants with significantly different predicted pHopt values compared to the wildtype, could be selected for site-saturated mutagenesis, aiding in the search for mutations that improve fitness at extreme pH levels. Third, integrating EpHod with existing protein design techniques could allow for large-scale screening of synthetically designed proteins and iterative selection of variants with improved functionality within specific pH ranges through machine-learning-directed evolution.^80–82^. Fourth, EpHod could be incorporated into conditional generative models to directly produce novel functional proteins with desired pH optima.^83,84^. Finally, the output of the final hidden layer of EpHod could serve as sequence embeddings that more richly capture pH optima, enabling subsequent supervised training (Supplementary Figure 12).

## Methods

### Data preparation

Enzyme optimum pH data were retrieved from BRENDA for 49,227 entries on May 25, 2022.^26^ Protein sequences could be downloaded from UniProt for 11,174 of these entries.^79^ Multiple pHopt entries for the same sequence were replaced with the average value, if the range was less than 1.0 pH unit, or else the sequence was discarded. To keep with the length limit of most protein language models, sequences with more than 1,022 residues or fewer than 32 residues were discarded, leaving 9,855 sequences that formed the pHopt dataset. A random selection of 1,971 sequences (20%) was set aside as the held-out testing set. The remaining sequences were clustered with mmseqs2 at 20% identity and the clusters were randomly split at 90% and 10% (7,124 and 760 sequences) for training and validation, respectively.^85^ The test set was further split into bins according to maximum sequence identity with the training set to enable analysis of the relationship between performance and sequence identity (Figure 3F and Supplementary Figure 8). Furthermore, to address concerns about some pHopt values being derived from a single pH value, or a relatively narrow range of pH measurements in experiments, those entries in the testing set that were annotated as “assay at” in BRENDA were removed to form a smaller “clean” testing set of 993 sequences as was done in a previous work.^49^ Regardless of the BRENDA annotation, we decided to use the full data for training and validation, since the models can potentially benefit from the relationship between the enzyme sequence and the pH range for activity. In evaluation, we found that EpHod exhibits similar performance on the full dataset and the “clean” dataset, suggesting that the model training was robust to the experiment noise in the dataset due to some pHopt values being derived from a limited a pH range (Supplementary Figure 13).

Optimum environment pH data for 5,621 bacteria were retrieved from BacDive on June 20, 2022 for the pHenv dataset.^29^ Multiple organismal entries were averaged if the range was less than 1.0 pH unit, or else they were discarded. Protein sequence data associated with each organism could be accessed for 3,715 organisms from NCBI RefSeq database and were downloaded using BioPython.^30,86^ Very short and long sequences were also discarded (<32 or >1022), resulting in 17.0 million sequences. Using SignalP 6.0,^31^ 2.2 million proteins were predicted to be secreted proteins possessing a signal peptide, and non-secreted proteins were discarded. All pHopt and pHenv sequences were combined and clustered with mmseqs2 at 20% sequence identity.^85^ Sequences in the pHenv training set that were grouped in the same cluster as any sequence in the pHopt testing set were discarded to prevent data leakage between the two datasets during transfer learning, leaving 1.9 million proteins in the pHenv dataset. The remaining pHenv clusters were randomly split into training and validation sets (90% and 10%) with 1.71 million and 188,525 sequences respectively. After clustering and splitting the datasets, there were 567 enzymes in the pHopt dataset that could be connected to 97 organisms with pHenv values, with a Pearson’s correlation coefficient of 0.17 between pHopt and pHenv values (Supplementary Figure 1).

### Loss reweighting

To mitigate the potentially adverse effects of the imbalance of pHopt label values,^36,37^ the loss function of each model was reweighted with values derived via five methods: *bin inverse, LDS inverse, bin sqrt, LDS sqrt*, and *LDS extreme* (Figure 1B). In the *bin inverse* method, the label values were categorized into three distinct bins, each associated with acidic, neutral, and alkaline pH ranges. These pH ranges are designated as *B*_*acidic*_ = {*y* | *y* ≤ 5}, *B*_*neutral*_ = {*y* |5 < *y* < 9}, and *B*_*alkaline*_ = {*y* | *y* ≥ 9}, respectively, where *y* is the pHopt label. The reciprocal of the size of each bin was used as the weight for all samples within the respective bin.

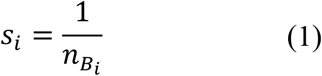

In the *LDS inverse* method, the reciprocal of label distribution smoothing (LDS) values were calculated according to Yang *et al*.^38^ First, the target values were split into 100 uniformly spaced bins to derive the empirical label distribution. Subsequently, a Gaussian kernel was applied to the binned values to estimate the effective label distribution via kernel density estimation (KDE). Hence, the assigned weight for each sample, *s*_*i*_, was computed as the reciprocal of the estimated density value, and is given by

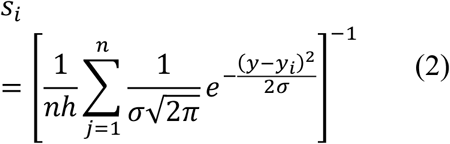

where *y*_*i*_ is the target value, *y* is the center value of the bin, *n* is the number of bins, and *h* and *σ* are the bandwidth and standard deviation of the Gaussian kernel that determine the smoothness and shape of the kernel, respectively. We used default values of 2 and 5 for *h* and *σ*, respectively.^38^ To attenuate the upweighting of rare acidic and alkaline samples, since inverse reweighting strategies may overcorrect the imbalance and lead to suboptimal performance,^87^ we examined sample weights derived by computing the square root of the *bin inverse* and *LDS inverse* weights, termed *bin sqrt* and *LDS sqrt*, respectively. This process yielded reweighting values that were intermediate between the inverse reweighting techniques and no reweighting (Figure 1B). In contrast, we evaluated a reweighting strategy (*LDS extreme*) that amplified the *LDS inverse* weights of scarce acidic and alkaline samples twofold, relative to abundant neutral values. The optimal reweighting techniques for each machine learning method were identified through hyperparameter optimization, employing either random search or grid search.^48^

For training neural networks, the root mean squared error (RMSE), reweighted by mean-normalized sample weights, was utilized as the loss function as follows:

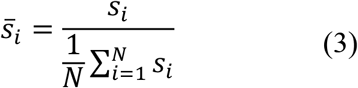

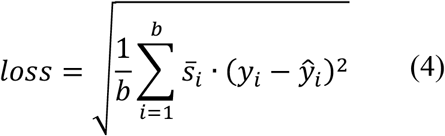

where N is the training data size and b is the size of each batch. For a given learning rate, we found that normalizing the sample weights helped prevent instability arising from high variance in gradient estimates.^88^ In place of loss reweighting for k-neighbor regression (KNR) models, we resampled the training data, drawing from the probability distribution defined by the sample weights.

### Sequence representation

Protein sequences were numerically represented using amino acid composition (AAC), one-hot encoding of the sequence (One-Hot), a collection of engineered features from the iFeature package (iFeature),^40^ and embeddings from six self-supervised protein language models (PLM) (Supplementary Table 2). The AAC for each protein was computed by determining the relative abundance of the canonical amino acids, yielding a 20-dimensional vector. One-hot encoding yielded binary representations of the sequences, with ones indicating the amino acid present at each residue, and zeros indicating all other amino acids. To achieve a uniform length, all sequences were post-padded with zeros up to 1024 residues, generating the representation, *x*_*onehot*_ ∈ ℝ^1024×20^, or the flattened vector, *x*_*onehot*_ ∈ ℝ^1×20480^. We derived 5,494 features from the iFeature package (Supplementary Table 7). For deriving PLM embeddings, the hidden states of the final transformer layer, or the final ByteNet convolution layer in CARP, were obtained. These per-residue embeddings, denoted as *x*_*emb*_ ∈ ℝ^*L*×*d*^, where *L* represents the sequence length and, *d* represents the hidden state dimension, were also averaged over the sequence length to obtain averaged embeddings, *x*_*avg*_ ∈ ℝ^1×*d*^. Embeddings from autoregressive models (ProGen2 and Tranception) were obtained by processing the sequences left-to-right (or N-to-C terminus) only. While a comprehensive exploration of other PLM embeddings for predicting pHopt could be beneficial, we focused on these six models to represent transformer- and convolution-based models, as well as masked and autoregressive models.

### Traditional machine learning methods

Traditional machine learning methods including, ridge regression (Ridge), random forest regression (RForest), support vector regression (SVR), k-neighbors regression (KNR), and XGBoost were implemented. Optimization of key hyperparameters was performed for each method, while default settings were employed for other hyperparameters. Ridge regression, a linear model, is inherently limited at capturing non-linear relationships between the features and target values. This is evident by the superior performance of non-linear models, which are better suited for capturing complex relationships in the data. Using the 5,494 hand-engineered features from iFeature, feature selection was performed to obtain an optimal subset of the features for predicting pHopt using principal components analysis (PCA) and recursive feature elimination (RFE).^89^ We chose RFE and PCA as feature selection approaches due to their efficiency and interpretability. With PCA, the top 1000 principal components were selected, explaining 80% of the data variance. With the RFE approach, a random forest with 20-trees was trained with all the features excluding AAC. The features were ranked based on the random forest Gini importance, and 10% of the features with the lowest Gini scores were discarded. The training process was repeated multiple times, discarding 10% of the features in each iteration and evaluating the RMSE, until one feature was left. The optimal feature combination corresponded to 33 features and the final random forest model was trained using 53 features (20 AAC and 33 RFE selection) (Supplementary Figures 14 and 15).

### Neural network methods

The initial architecture of the neural network models was based on notable models from the literature.^28,71,83,90^ However, during hyperparameter optimization, we explored variations in the architecture to identify more optimal configurations for this task (Supplementary Table 4). Feed-forward neural networks (FNN) contained up to three hidden dense layers with the same number of units in each layer, ranging from 128 to 1,024. Batch normalization was applied to the output of the dense layers, followed by a non-linear activation and dropout. An optional residual (skip) connection was employed, connecting the output of the first and last hidden layers. Convolutional neural networks (CNN) consisted of up to 8 blocks of repeating convolutional, batch normalization, non-linear activation, and pooling layers. The initial convolutional layer filter size was set at 32, and successively doubled for each subsequent layer. Pooling layers utilized a kernel size of 2 and a stride of 2. All input tensors to the convolution layers were ‘same’ padded. The output from the last convolution block was flattened and supplied to a feed-forward network to generate the final pHopt prediction. Dilated convolution neural networks (DCNN) contained up to 6 blocks of 10 convolution layers with exponentially increasing dilation rates from 1, 2, 4, up to 512, and a kernel size of 2.^90,91^ The input tensors to the convolution layers were causal padded to ensure unidirectional processing from N to C terminus. Each convolution layer was followed by batch normalization and non-linear activation layers and a dropout layer was applied after each block. The output from the last block was passed to a feed-forward neural network to predict target values. Recurrent neural networks (RNN) employed a unidirectional gated recurrent unit (GRU) processing the sequence tensors from N-to-C terminus.^92^ The final hidden state of the GRU, corresponding to the C-terminal residue, was connected to a feed-forward network with one hidden layer to predict target values. It is well known that the problem of vanishing and exploding gradients pose challenges in effectively training RNNs.^93^ To address these challenges, the norm of the gradients was clipped to 1.0. However, after hyperparameter optimization, the RNN consistently underperformed all other neural network methods, including FNN, CNN, and DCNN, likely due to the vanishing gradient problem, at least in part. We also trained a long short-term memory (LSTM) model with optimal hyperparameters of the GRU,^94^ but this did not lead to performance gains compared with the GRU. While truncated backpropagation through time (TBPTT) may help alleviate vanishing gradients,^95^ evaluation of RNNs with TBPTT was beyond the scope of this work. In any case, our results highlight the difficulty of training RNNs on biological sequences as observed previously,^90^ and demonstrate that dilated convolutions may be more suitable for sequence modeling of proteins.^46^

We hypothesized that valuable information is lost in averaged embeddings (average pooling) regarding the relative importance of specific residues that could improve predictive performance. Hence, we trained neural networks on full per-residue embeddings. With pooling, each residue’s embedding contributes equally to the averaged embedding such that the averaged embeddings can be expressed as a weighted sum of the per-residue embeddings with equal weights of 1/*L* as follows,

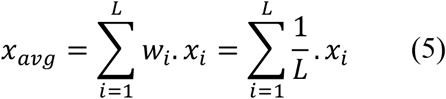

Alternatively, the weights, *w*_*i*_, can be variably learned to capture the relative importance of each residue, *i*. Following the light attention (LAT) module proposed by Stärk *et al*.,^47^ in place of equal weights in the case of average pooling, we fit two separate convolutions with a specified filter size and ‘same’ padding on full per-residue embeddings, *x*_*emb*_ ∈ ℝ^*L*×*d*^, to derive learned attention weights and transformed values, *x*_*attn*_ ∈ ℝ^*L*×*d*^ and *x*_*val*_ ∈ ℝ^*L*×*d*^, respectively. The attention coefficients were softmax-normalized over the length dimension to obtain attention weights per residue that sum to 1.0. Consequently, the attention-weighted representation, *x*_*sum*_ ∈ ℝ^1×*d*^, was obtained for each protein by summing the element-wise product of *x*_*attn*_ and *x*_*val*_ over the sequence dimension as follows,

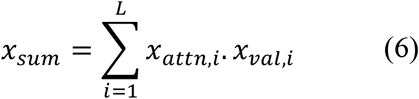

Comparing (5) and (6), *x*_*attn*_ enables the LAT model to learn varying weights for each residue, rather than 1/*L* in averaging, and potentially captures the specific contribution of each residue. Stärk *et al*. also showed that, compared to using *x*_*sum*_ alone as the protein representation, concatenating *x*_*sum*_ with the representation derived from max-pooling *x*_*val*_ over the sequence length to yield *x*_*rep*_ ∈ ℝ^*L*×2*d*^, led to performance gains in predicting protein localization.^47^ Hence, we similarly concatenated attention-weighted values with max-pooled values in our implementation of the LAT architecture (Figure 3A).

We also proposed variations to the LAT architecture for possible improvements in performance. We experimented with perceptive light attention (PAT), in which the average embedding, *x*_*avg*_ ∈ ℝ^1×*d*^, is repeated *L* times and concatenated with the ESM-1v embedding tensor, *x*_*emb*_ ∈ ℝ^*L*×*d*^, to obtain a perceptive embedding, *x*_*perc*_ ∈ ℝ^2*L*×*d*^, from which we derive attention weights and values, rather than from *x*_*emb*_ as in the LAT model. The motivation is to expand the perceptive breadth of the convolution layers beyond the kernel size, which does not consider all residues in each stride. We also proposed dilated convolution with LAT (DCAT) which incorporates a single residual block of a DCNN with 9 convolution layers that is fit on the element-wise product of the attention weights and values, *x*_*attn*_ and *x*_*val*_, respectively. The output of the dilated convolution block is subsequently flattened and passed to a feed-forward network to predict target values. We evaluated residual light attention (RLAT), with multiple residual dense layers replacing the single hidden layer in LAT. We anticipated that this modification might provide improvements by leveraging deeper layers to better capture the nonlinear mapping between the embeddings and target values. We also examined the performance of RNNs (GRU), processing the embeddings in the N-to-C direction, and feeding the final RNN hidden states to a single hidden layer FNN. After extensive hyperparameter optimization and evaluation, the PAT, DCAT, and RNN methods underperformed LAT, while RLAT slightly outperformed LAT. Based on this, we further trained the RLAT model with optimal hyperparameters using a transfer learning approach, by first pre-training on 1.7 million pHenv values and fine-tuning on 7,124 pHopt values.

### Training routine

Traditional machine learning methods were implemented with the scikit-learn Python package, (version 0.23.2),^96^ and neural network models were built with PyTorch.^97^ Training of neural networks was carried out on the Eagle high-performance computing cluster (HPC) at the National Renewable Energy Laboratory, USA, with 16 GB NVIDIA Tesla V100 PCle GPUs. The Adam optimization algorithm was applied with default settings, and the initial learning rate was optimized for each method. To prevent overfitting and ensure efficient training, a combination of dropout, L2 regularization, batch normalization, and early stopping were implemented. A learning rate schedule was applied that lowered the learning rate by a factor of 0.5, if the validation loss did not improve after 10 epochs, and stopped the training if the validation loss did not improve after 20 epochs. A batch size of 32 was employed throughout. In the unique case of pre-training the RLAT model on the pHenv dataset (Figure 3D), an initial learning rate of 10^−3^ was used since this was the optimal learning rate in hyperparameter tuning of the RLAT model. This learning rate was reduced if the validation loss did not improve after one epoch, and the training was stopped if validation loss did not improve after five epochs.

### Performance evaluation

Regression performance was accessed using RMSE, R^2^, and Pearson and Spearman correlation coefficients. Given the considerable variability in the scale and precision of pHopt measurements in experiments, the ability of the models to correctly classify the pHopt range (as acidic, neutral, or alkaline) was additionally evaluated by computing the percent accuracy, Matthew’s correlation coefficient (MCC), F1 score, and the area under the receiver operator characteristic curve (AUC). To compute the F1 and AUC scores, which are typically binary metrics, the performance was evaluated for each class compared with the other two classes, and the scores for all three classes were averaged. All performance metrics were computed utilizing the scikit-learn Python package. Since the pH data was imbalanced, with most pH values falling within the range of 5 to 9, it was important to address this imbalance in both training and evaluation, to ensure that the trained models accurately predict the pHopt beyond the over-represented neutral pH range, and that the estimated performance effectively represents the entire pH range. Hence, sample-weights (*bin-inverse*) were applied in evaluation for all performance metrics. For the evaluation of correlation coefficients, the testing dataset was resampled by drawing from the distribution defined by sample weights, to achieve a balanced testing dataset, and averaged over 100 resampling repetitions.^36^

### Interpretability analyses and structural evaluation

To assess what EpHod learned in training on a per-residue basis, the attention weights generated by the softmax layer of the RLAT module were evaluated. The weights were averaged over the feature dimension and normalized by dividing with the mean value. To understand important residues EpHod focused on for prediction, the normalized attention weights for each residue were compared with the amino acid type, the distance from the active site, and the solvent accessible surface area (SASA). Three-dimensional structures for enzymes in the pHopt dataset were retrieved from the AlphaFold protein structure database (AlphaFoldDB).^98^ For consistency, AlphaFold structures were utilized for all structural analyses, even when a crystal structure was available. Enzymes lacking a structure in AlphaFoldDB were folded with ColabFold.^99^ To compare per-residue attention weights with distance to the active site, 3,074 enzymes in the dataset which had at least one residue annotated as “Active Site” in UniProt were considered. In the AlphaFold structure, the distance to the active site for each residue was computed as the Euclidean distance between the alpha carbon of the residue and the centroid of the alpha carbons of all annotated active-site residues. SASA calculations were performed on the AlphaFold structures using the Shrake and Rupley algorithm with default parameters via the Biopython module.^86,100^ To distinguish surface or exposed residues from buried residues, the relative SASA was computed by dividing the SASA by the maximum SASA from the Sander and Rost scale,^101^ and a residue was classified as buried if the relative SASA was less than 20%.^102^ The relative composition of positively to negatively charged amino acids on the protein surface (RKDE) was computed by evaluating the amino acid composition of residues among surface residues, and computing, *c*_*Arg*_ + *c*_*Lys*_ ™ *c*_*Asp*_ ™ *c*_*Glu*_, where *c*_*a*_ is the percent composition of amino acid, *a*, on the protein surface. The pH for minimum (optimum) free energy of folding, ΔΔG pHopt, and the isoelectric point, pI, were computed using PROPKA3 with default settings on the AlphaFold structures.^65^

## Data availability

Data for training the models, as well as the weights of the optimal EpHod model, are available at https://doi.org/10.5281/zenodo.8011248.

## Code availability

Code for EpHod and training the other models is available at https://github.com/beckham-lab/EpHod.

## Acknowledgements

We thank Ralph Estanboulieh and Rose Orenbuch for their support in visualizations and analyses. We thank Jeffery Law, Aaron Kollasch, Daniel Ritter, Pascal Notin, Jonathan Frazer, and Malfalda Dias for helpful discussions. This material is based upon work supported by the U.S. Department of Energy, Office of Science, Office of Biological and Environmental Research, Genomic Science Program under Award Number DE-SC0022024 to NPG. This work was authored in part by Alliance for Sustainable Energy, LLC, the manager and operator of the National Renewable Energy Laboratory for the U.S. Department of Energy (DOE) under Contract No. DE-AC36-08GO28308. Partial funding to JEG and GTB was provided by the U.S. Department of Energy, Office of Energy Efficiency and Renewable Energy, Advanced Materials and Manufacturing Technologies Office (AMMTO) and Bioenergy Technologies Office (BETO) as part of the BOTTLE™ Consortium and was supported by AMMTO and BETO under contract no. DE-AC36-08GO28308 with the National Renewable Energy Laboratory, operated by Alliance for Sustainable Energy, LLC. Partial funding to JEG and GTB was also provided by the U.S. Department of Energy Office of Energy Efficiency and Renewable Energy Bioenergy Technologies Office (BETO) for the Agile BioFoundry. We thank Gayle Bentley at DOE and members of the Agile BioFoundry for helpful discussions. Partial funding to JEG and GTB was also provided by the U.S. Department of Energy Office of Science Biological and Environmental Research via DE-SC0023278. The views expressed herein do not necessarily represent the views of the DOE or the U.S. Government. The U.S. Government retains, and the publisher, by accepting the article for publication, acknowledges that the U.S. Government retains a non-exclusive, paid-up, irrevocable, worldwide license to publish or reproduce the published form of this work, or allow others to do so, for U.S. Government purposes.

## Author contributions

GTB and JEG conceived of the project, and JEG, DM, NPG, CS, and GTB designed the study. JEG trained the predictive models, and AYS, MK, and JEG evaluated the models. The manuscript was written by JEG and edited and approved by all authors.

## References

1 Barroca, M. et al. Deciphering the factors defining the pH-dependence of a commercial glycoside hydrolase family 8 enzyme. Enzyme Microb. Technol. 96, 163–169 (2017).

2 Reed, C. J., Lewis, H., Trejo, E., Winston, V. & Evilia, C. Protein adaptations in archaeal extremophiles. Archaea 2013, e373275 (2013).

3 Protze, J. et al. An extracellular tetrathionate hydrolase from the thermoacidophilic archaeon Acidianus ambivalens with an activity optimum at pH 1. Front. Microbiol. 2, (2011).

4 Pradeep, G. C. et al. An extremely alkaline novel chitinase from Streptomyces sp. CS495. Process Biochem. 49, 223–229 (2014).

5 Ferrer, M., Golyshina, O., Beloqui, A. & Golyshin, P. N. Mining enzymes from extreme environments. Curr. Opin. Microbiol. 10, 207–214 (2007).

6 Verma, D. & Satyanarayana, T. Xylanolytic extremozymes retrieved from environmental metagenomes: Characteristics, genetic engineering, and applications. Front. Microbiol. 11, (2020).

7 Shahraki, M. F. et al. A computational learning paradigm to targeted discovery of biocatalysts from metagenomic data: A case study of lipase identification. Biotechnol. Bioeng. 119, 1115–1128 (2022).

8 Erickson, E. et al. Sourcing thermotolerant poly(ethylene terephthalate) hydrolase scaffolds from natural diversity. Nat. Commun. 13, 7850 (2022).

9 Wang, C.-H., Liu, X.-L., Huang, R.-B.He, B.-F. & Zhao, M.-M. Enhanced acidic adaptation of Bacillus subtilis Ca-independent alpha-amylase by rational engineering of pKa values. Biochem. Eng. J. 139, 146–153 (2018).

10 dos Santos, J. P., Zavareze, E. da R., Dias, A. R. G. & Vanier, N. L. Immobilization of xylanase and xylanase–β-cyclodextrin complex in polyvinyl alcohol via electrospinning improves enzyme activity at a wide pH and temperature range. Int. J. Biol. Macromol. 118, 1676–1684 (2018).

11 Giri, P., Pagar, A. D., Patil, M. D. & Yun, H. Chemical modification of enzymes to improve biocatalytic performance. Biotechnol. Adv. 53, 107868 (2021).

12 Xue, Y. et al. Chemical modification of stem bromelain with anhydride groups to enhance its stability and catalytic activity. J. Mol. Catal. B Enzym. 63, 188–193 (2010).

13 Li, C. Effects of chemical modification by chitooligosaccharide on enzyme activity and stability of yeast β-D-fructofuranosidase. Enzyme Microb. Technol. 64–65, 24–32 (2014).

14 Li, S.-F., Cheng, F.Wang, Y.-J. & Zheng, Y.-G. Strategies for tailoring pH performances of glycoside hydrolases. Crit. Rev. Biotechnol. 43, 121–141 (2023).

15 Shi, X., Wu, D., Xu, Y. & Yu, X. Engineering the optimum pH of β-galactosidase from Aspergillus oryzae for efficient hydrolysis of lactose. J. Dairy Sci. 105, 4772–4782 (2022).

16 Hebditch, M. & Warwicker, J. Web-based display of protein surface and pH-dependent properties for assessing the developability of biotherapeutics. Sci. Rep. 9, 1969 (2019).

17 Schmitz, M. et al. patcHwork: a user-friendly pH sensitivity analysis web server for protein sequences and structures. Nucleic Acids Res. 50, W560–W567 (2022).

18 Oeller, M. et al. Sequence-based prediction of pH-dependent protein solubility using CamSol. Brief. Bioinform. 24, bbad004 (2023).

19 Zhang, G., Li, H. & Fang, B. Discriminating acidic and alkaline enzymes using a random forest model with secondary structure amino acid composition. Process Biochem. 44, 654–660 (2009).

20 Lin, H., Chen, W. & Ding, H. AcalPred: A sequence-based tool for discriminating between acidic and alkaline enzymes. PLOS ONE 8, e75726 (2013).

21 Fan, G.-L.Li, Q.-Z. & Zuo, Y.-C. Predicting acidic and alkaline enzymes by incorporating the average chemical shift and gene ontology informations into the general form of Chou’s PseAAC. Process Biochem. 48, 1048–1053 (2013).

22 Khan, Z. U., Hayat, M. & Khan, M. A. Discrimination of acidic and alkaline enzyme using Chou’s pseudo amino acid composition in conjunction with probabilistic neural network model. J. Theor. Biol. 365, 197–203 (2015).

23 Yan, S. & Wu, G. Predicting pH optimum for activity of beta-glucosidases. J. Biomed. Sci. Eng. 12, 354–367 (2019).

24 Wang, X., Li, H., Gao, P., Liu, Y. & Zeng, W. Combining support vector machine with dual g-gap dipeptides to discriminate between acidic and alkaline enzymes. Lett. Org. Chem. 16, 325–331 (2019).

25 Li, X. et al. A sequence embedding method for enzyme optimal condition analysis. BMC Bioinformatics 21, 512 (2020).

26 Schomburg, I. et al. The BRENDA enzyme information system–From a database to an expert system. J. Biotechnol. 261, 194–206 (2017).

27 Puissant, J. et al. The pH optimum of soil exoenzymes adapt to long term changes in soil pH. Soil Biol. Biochem. 138, 107601 (2019).

28 Li, G. et al. Learning deep representations of enzyme thermal adaptation. Protein Sci. 31, e4480 (2022).

29 Reimer, L. C. et al. BacDive in 2022: the knowledge base for standardized bacterial and archaeal data. Nucleic Acids Res. 50, D741–D746 (2022).

30 Sayers, E. W. et al. Database resources of the National Center for Biotechnology Information. Nucleic Acids Res. 49, D10–D17 (2021).

31 Teufel, F. et al. SignalP 6.0 predicts all five types of signal peptides using protein language models. Nat. Biotechnol. 40, 1023–1025 (2022).

32 Booth, I. R. Regulation of cytoplasmic pH in bacteria. Microbiol. Rev. 49, 359–378 (1985).

33 Baker-Austin, C. & Dopson, M. Life in acid: pH homeostasis in acidophiles. Trends Microbiol. 15, 165–171 (2007).

34 Hough, D. W. & Danson, M. J. Extremozymes. Curr. Opin. Chem. Biol. 3, 39–46 (1999).

35 Tan, C. et al. A Survey on deep transfer learning. Preprint at 10.48550/arXiv.1808.01974 (2018).

36 Gado, J. E., Beckham, G. T. & Payne, C. M. Improving enzyme optimum temperature prediction with resampling strategies and ensemble learning. J. Chem. Inf. Model. 60, 4098–4107 (2020).

37 Branco, P., Torgo, L. & Ribeiro, R. P. Pre-processing approaches for imbalanced distributions in regression. Neurocomputing 343, 76–99 (2019).

38 Yang, Y., Zha, K., Chen, Y.-C., Wang, H. & Katabi, D. Delving into deep imbalanced regression. Preprint at 10.48550/arXiv.2102.09554 (2021).

39 Meier, J. et al. Language models enable zero-shot prediction of the effects of mutations on protein function. 2021.07.09.450648 Preprint at 10.1101/2021.07.09.450648 (2021).

40 Chen, Z. et al. iFeature: a Python package and web server for features extraction and selection from protein and peptide sequences. Bioinforma. Oxf. Engl. 34, 2499–2502 (2018).

41 Unsal, S. et al. Learning functional properties of proteins with language models. Nat. Mach. Intell. 4, 227–245 (2022).

42 Rives, A. et al. Biological structure and function emerge from scaling unsupervised learning to 250 million protein sequences. Proc. Natl. Acad. Sci. 118, e2016239118 (2021).

43 Elnaggar, A. et al. ProtTrans: toward understanding the language of life through self-supervised learning. IEEE Trans. Pattern Anal. Mach. Intell. 44, 7112–7127 (2022).

44 Notin, P. et al. Tranception: protein fitness prediction with autoregressive transformers and inference-time retrieval. Preprint at 10.48550/arXiv.2205.13760 (2022).

45 Nijkamp, E., Ruffolo, J., Weinstein, E. N., Naik, N. & Madani, A. ProGen2: exploring the boundaries of protein language models. Preprint at 10.48550/arXiv.2206.13517 (2022).

46 Yang, K. K., Lu, A. X. & Fusi, N. Convolutions are competitive with transformers for protein sequence pretraining. 2022.05.19.492714 Preprint at 10.1101/2022.05.19.492714 (2022).

47 Stärk, H., Dallago, C., Heinzinger, M. & Rost, B. Light attention predicts protein location from the language of life. Bioinforma. Adv. 1, vbab035 (2021).

48 Bergstra, J. & Bengio, Y. Random search for hyper-parameter optimization. J. Mach. Learn. Res. 13, 281–305 (2012).

49 Li, G. et al. Performance of Regression Models as a Function of Experiment Noise. Bioinforma. Biol. Insights 15, 11779322211020315 (2021).

50 Detlefsen, N. S., Hauberg, S. & Boomsma, W. Learning meaningful representations of protein sequences. Nat. Commun. 13, 1914 (2022).

51 Zaretckii, M., Buslaev, P., Kozlovskii, I., Morozov, A. & Popov, P. Approaching Optimal pH Enzyme Prediction with Large Language Models. ACS Synth. Biol. (2024) doi:10.1021/acssynbio.4c00465.

52 Kroll, A. & Lercher, M. J. Machine learning models for the prediction of enzyme properties should be tested on proteins not used for model training. 2023.02.06.526991 Preprint at 10.1101/2023.02.06.526991 (2023).

53 Suplatov, D. et al. Computational design of a pH stable enzyme: understanding molecular mechanism of penicillin acylase’s adaptation to alkaline conditions. PLOS ONE 9, e100643 (2014).

54 Huang, Y., Krauss, G., Cottaz, S., Driguez, H. & Lipps, G. A highly acid-stable and thermostable endo-beta-glucanase from the thermoacidophilic archaeon Sulfolobus solfataricus. Biochem. J. 385, 581–588 (2005).

55 Mamo, G., Thunnissen, M., Hatti-Kaul, R. & Mattiasson, B. An alkaline active xylanase: insights into mechanisms of high pH catalytic adaptation. Biochimie 91, 1187–1196 (2009).

56 Wang, Y., Xu, M., Yang, T., Zhang, X. & Rao, Z. Surface charge-based rational design of aspartase modifies the optimal pH for efficient β-aminobutyric acid production. Int. J. Biol. Macromol. 164, 4165– 4172 (2020).

57 Jakob, F. et al. Surface charge engineering of a Bacillus gibsonii subtilisin protease. Appl. Microbiol. Biotechnol. 97, 6793–6802 (2013).

58 Yang, T. et al. N20D/N116E combined mutant downward shifted the pH optimum of Bacillus subtilis NADH Oxidase. Biology 12, 522 (2023).

59 Masui, A., Fujiwara, N., Yamamoto, K., Takagi, M. & Imanaka, T. Rational design for stabilization and optimum pH shift of serine protease AprN. J. Ferment. Bioeng. 85, 30–36 (1998).

60 Turunen, O., Vuorio, M., Fenel, F. & Leisola, M. Engineering of multiple arginines into the Ser/Thr surface of Trichoderma reesei endo-1,4-β-xylanase II increases the thermotolerance and shifts the pH optimum towards alkaline pH. Protein Eng. Des. Sel. 15, 141–145 (2002).

61 Li, Q., Jiang, T., Liu, R., Feng, X. & Li, C. Tuning the pH profile of β-glucuronidase by rational site-directed mutagenesis for efficient transformation of glycyrrhizin. Appl. Microbiol. Biotechnol. 103, 4813–4823 (2019).

62 Pokhrel, S., Joo, J. C. & Yoo, Y. J. Shifting the optimum pH of Bacillus circulans xylanase towards acidic side by introducing arginine. Biotechnol. Bioprocess Eng. 18, 35–42 (2013).

63 Carvalho, D. V., Pereira, E. M. & Cardoso, J. S. Machine Learning Interpretability: A Survey on Methods and Metrics. Electronics 8, 832 (2019).

64 Austin, H. P. et al. Characterization and engineering of a plastic-degrading aromatic polyesterase. Proc. Natl. Acad. Sci. U. S. A. 115, E4350–E4357 (2018).

65 Olsson, M. H. M., Søndergaard, C. R., Rostkowski, M. & Jensen, J. H. PROPKA3: Consistent treatment of internal and surface residues in empirical pKa predictions. J. Chem. Theory Comput. 7, 525–537 (2011).

66 Talley, K. & Alexov, E. On the pH-optimum of activity and stability of proteins. Proteins 78, 2699– 2706 (2010).

67 Alexov, E. Numerical calculations of the pH of maximal protein stability. The effect of the sequence composition and three-dimensional structure. Eur. J. Biochem. 271, 173–185 (2004).

68 Yu, T. et al. Enzyme function prediction using contrastive learning. Science 379, 1358–1363 (2023).

69 Pak, M. A., Dovidchenko, N. V., Sharma, S. M. & Ivankov, D. N. New mega dataset combined with deep neural network makes a progress in predicting impact of mutation on protein stability. 2022.12.31.522396 Preprint at 10.1101/2022.12.31.522396 (2023).

70 Kroll, A., Rousset, Y., Hu, X.-P., Liebrand, N. A. & Lercher, M. J. Turnover number predictions for kinetically uncharacterized enzymes using machine and deep learning. Nat. Commun. 14, 4139 (2023).

71 Gelman, S., Fahlberg, S. A., Heinzelman, P., Romero, P. A. & Gitter, A. Neural networks to learn protein sequence–function relationships from deep mutational scanning data. Proc. Natl. Acad. Sci. 118, e2104878118 (2021).

72 Dallago, C. et al. FLIP: Benchmark tasks in fitness landscape inference for proteins. 2021.11.09.467890 Preprint at 10.1101/2021.11.09.467890 (2022).

73 Sledzieski, S. et al. Democratizing protein language models with parameter-efficient fine-tuning. Proc. Natl. Acad. Sci. 121, e2405840121 (2024).

74 Xu, M. et al. PEER: A Comprehensive and Multi-Task Benchmark for Protein Sequence Understanding. Adv. Neural Inf. Process. Syst. 35, 35156–35173 (2022).

75 Ferdous, S., Shihab, I. F. & Reuel, N. F. Effects of sequence features on machine-learned enzyme classification fidelity. Biochem. Eng. J. 187, 108612 (2022).

76 Li, G., Rabe, K. S., Nielsen, J. & Engqvist, M. K. M. Machine Learning Applied to Predicting Microorganism Growth Temperatures and Enzyme Catalytic Optima. ACS Synth. Biol. 8, 1411–1420 (2019).

77 Liu, H., HaoChen, J. Z., Gaidon, A. & Ma, T. Self-supervised learning is more robust to dataset imbalance. Preprint at 10.48550/arXiv.2110.05025 (2022).

78 Richardson, L. et al. MGnify: the microbiome sequence data analysis resource in 2023. Nucleic Acids Res. 51, D753–D759 (2023).

79 UniProt Consortium. UniProt: the Universal Protein Knowledgebase in 2023. Nucleic Acids Res. 51, D523–D531 (2023).

80 Hopf, T. A. et al. Mutation effects predicted from sequence co-variation. Nat. Biotechnol. 35, 128– 135 (2017).

81 Hie, B. L. & Yang, K. K. Adaptive machine learning for protein engineering. Curr. Opin. Struct. Biol. 72, 145–152 (2022).

82 Yang, K. K., Wu, Z. & Arnold, F. H. Machine-learning-guided directed evolution for protein engineering. Nat. Methods 16, 687–694 (2019).

83 Hawkins-Hooker, A. et al. Generating functional protein variants with variational autoencoders. PLoS Comput. Biol. 17, e1008736 (2021).

84 Strokach, A. & Kim, P. M. Deep generative modeling for protein design. Curr. Opin. Struct. Biol. 72, 226–236 (2022).

85 Steinegger, M. & Söding, J. MMseqs2 enables sensitive protein sequence searching for the analysis of massive data sets. Nat. Biotechnol. 35, 1026–1028 (2017).

86 Cock, P. J. A. et al. Biopython: freely available Python tools for computational molecular biology and bioinformatics. Bioinformatics 25, 1422–1423 (2009).

87 Cui, Y., Jia, M., Lin, T.-Y., Song, Y. & Belongie, S. Class-balanced loss based on effective number of samples. in 9268–9277 (2019).

88 Riesselman, A. J., Ingraham, J. B. & Marks, D. S. Deep generative models of genetic variation capture the effects of mutations. Nat. Methods 15, 816–822 (2018).

89 Guyon, I., Weston, J., Barnhill, S. & Vapnik, V. Gene selection for cancer classification using support vector machines. Mach. Learn. 46, 389–422 (2002).

90 Shin, J.-E. et al. Protein design and variant prediction using autoregressive generative models. Nat. Commun. 12, 2403 (2021).

91 Oord, A. van den et al. WaveNet: a generative model for raw audio. Preprint at 10.48550/arXiv.1609.03499 (2016).

92 Cho, K. et al. Learning phrase representations using RNN encoder–decoder for statistical machine translation. in Proceedings of the 2014 Conference on Empirical Methods in Natural Language Processing (EMNLP) 1724–1734 (Association for Computational Linguistics, Doha, Qatar, 2014). doi:10.3115/v1/D14-1179.

93 Pascanu, R., Mikolov, T. & Bengio, Y. On the difficulty of training recurrent neural networks. in Proceedings of the 30th International Conference on International Conference on Machine Learning - Volume 28 III-1310-III–1318 (JMLR.org, Atlanta, GA, USA, 2013).

94 Hochreiter, S. & Schmidhuber, J. Long short-term memory. Neural Comput. 9, 1735–1780 (1997).

95 Alley, E. C., Khimulya, G., Biswas, S., AlQuraishi, M. & Church, G. M. Unified rational protein engineering with sequence-based deep representation learning. Nat. Methods 16, 1315–1322 (2019).

96 Pedregosa, F. et al. Scikit-learn: machine learning in Python. J. Mach. Learn. Res. 12, 2825–2830 (2011).

97 Paszke, A. et al. PyTorch: an imperative Style, high-performance deep learning library. In Advances in Neural Information Processing Systems vol. 32 (Curran Associates, Inc., 2019).

98 Varadi, M. et al. AlphaFold Protein Structure Database: massively expanding the structural coverage of protein-sequence space with high-accuracy models. Nucleic Acids Res. 50, D439–D444 (2022).

99 Mirdita, M. et al. ColabFold: making protein folding accessible to all. Nat. Methods 19, 679–682 (2022).

100 Shrake, A. & Rupley, J. A. Environment and exposure to solvent of protein atoms. Lysozyme and insulin. J. Mol. Biol. 79, 351–371 (1973).

101 Rost, B. & Sander, C. Conservation and prediction of solvent accessibility in protein families. Proteins 20, 216–226 (1994).

102 Savojardo, C., Manfredi, M., Martelli, P. L. & Casadio, R. Solvent accessibility of residues undergoing pathogenic variations in humans: from protein structures to protein sequences. Front. Mol. Biosci. 7, (2021).

