## Supplemental Information for "Machine learning prediction of enzyme optimum pH"

### Table of Contents

#### Supplementary Figures


1 Relationship between the pHenv and pHopt datasets
2 Comparison of language model embeddings on validation set
3 Sample-weighted RMSE of pHopt values
4 Sample-weighted AUC of pHopt values
5 Sample-weighted MCC of pHopt values
6 Sample-weighted F1 score of pHopt values
7 Comparison of EpHod with OphPred.
8 Performance of the models after applying maximum sequence identity thresholds.
9 Average attention weights of residues in the pHopt dataset
10 Average composition of residues in the pHopt dataset
11 RMSE of ESM1v-RLAT on testing set split into subgroups
12 Comparison of EpHod and ESM-1v embeddings for supervised prediction tasks
13 Performance of EpHod on the full testing set and on a "clean" subset


#### Supplementary Tables


1 Taxonomical distribution of enzymes in the pHopt dataset
2 Protein language models used to derive embeddings for predicting pHopt values
3 Hyperparameter space of traditional machine learning models
4 Hyperparameter space of neural network models
5 Number of models trained with per-protein vector representations
6 Number of neural network models trained with per-residue representations
7 Description of 5,494 features generated by the iFeature

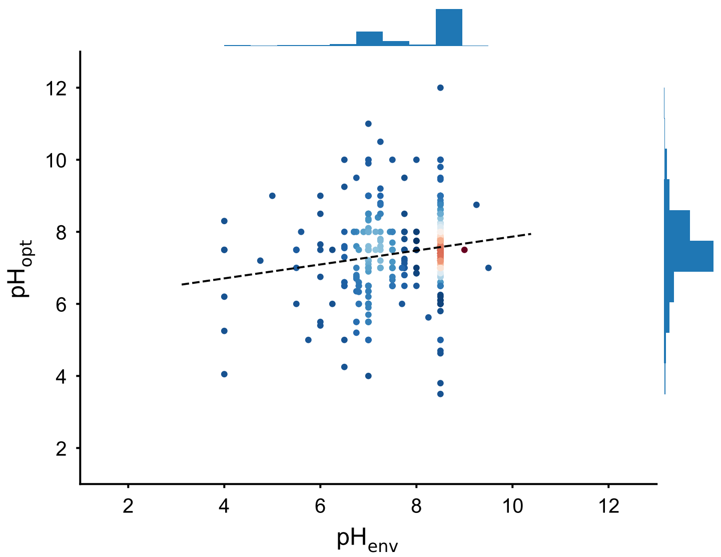


**Supplementary Figure 1**. Relationship between pHenv and pHopt values. From the 9,855 sequences in the pHopt dataset, 567 could be linked to 97 organisms having environment pH data in the pHenv dataset. The figure here shows moderate correlation between pHopt and pHenv for these 567 enzymes (Pearson R = 0.170, p-value = 4.8e-5; Spearman R = 0.155, p-value = 2.0e-4).

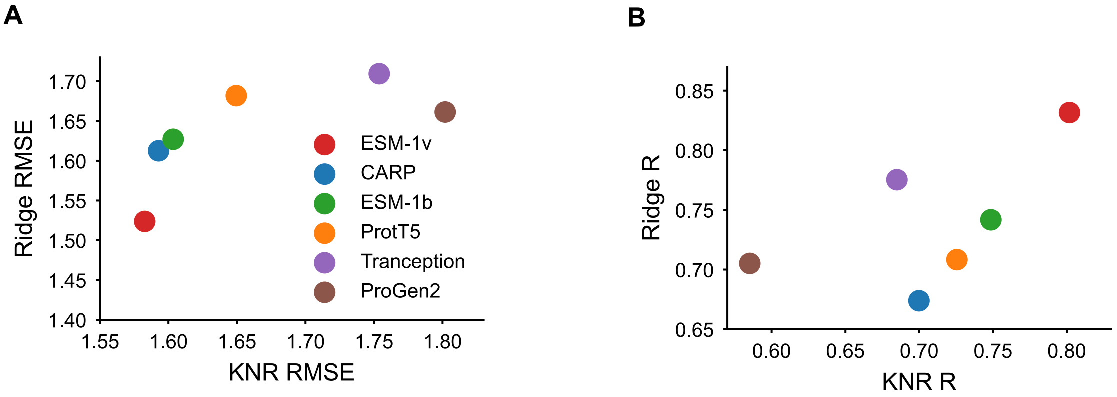


**Supplementary Figure 2.** Comparison of language model embeddings on validation set. **(A)** Root mean squared error (RMSE) of k-neighbor regression (KNR) and ridge regression models trained on averaged embeddings from six protein language models to predict the pHopt values of the validation set (n=760). **(B)** Pearson's correlation coefficient between predicted and experiment pHopt as in **A**. ESM-1v yielded the lowest error and highest correlation with both KNR and ridge models, and was subsequently selected for training with deep neural networks.

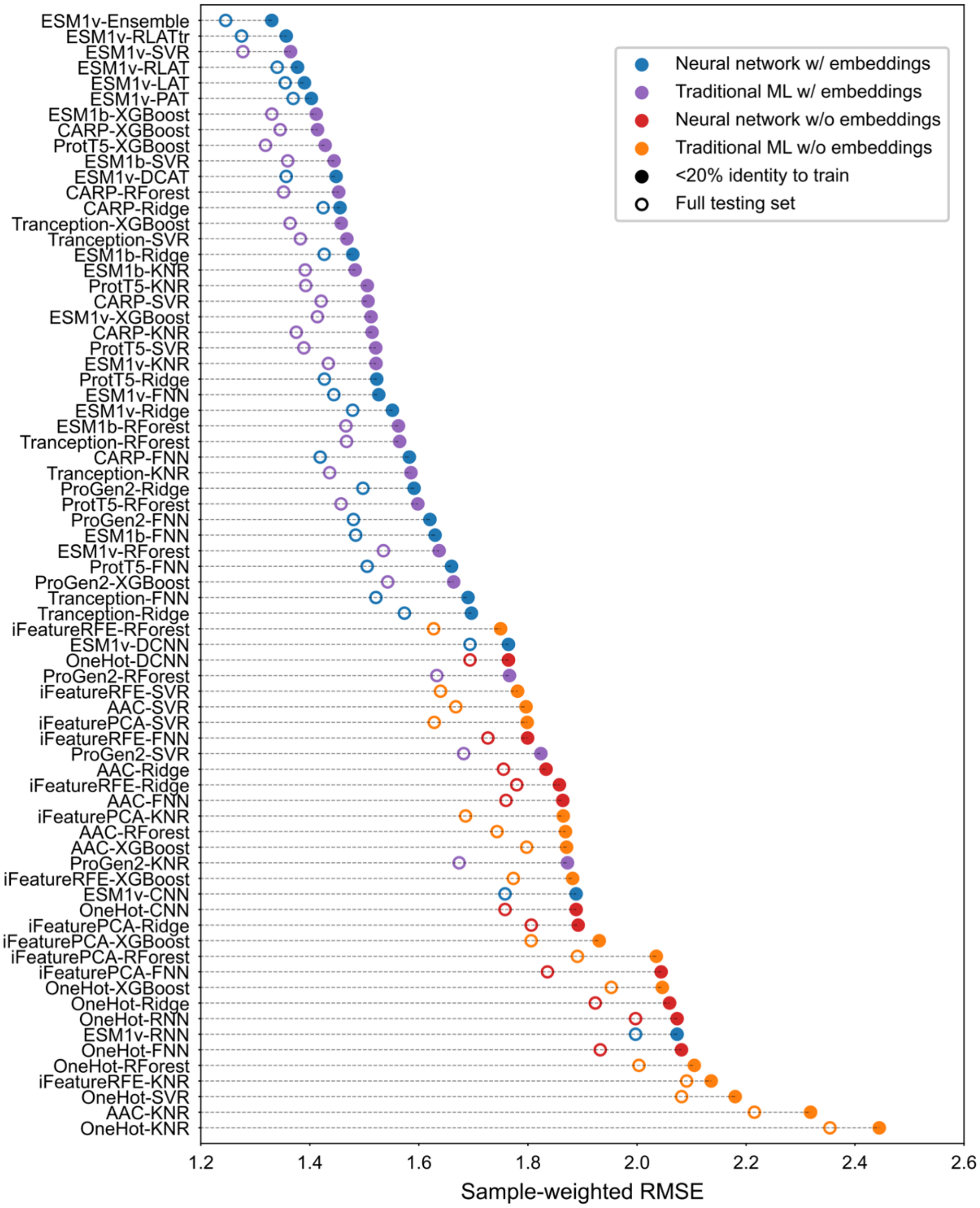


**Supplementary Figure 3.** Sample-weighted root mean squared error (RMSE) of pHopt values predicted by optimal models, evaluated on both the complete held-out pHopt testing set (n=1,971) and a subset of the testing set with less than 20% sequence identity to the training set (n=999).

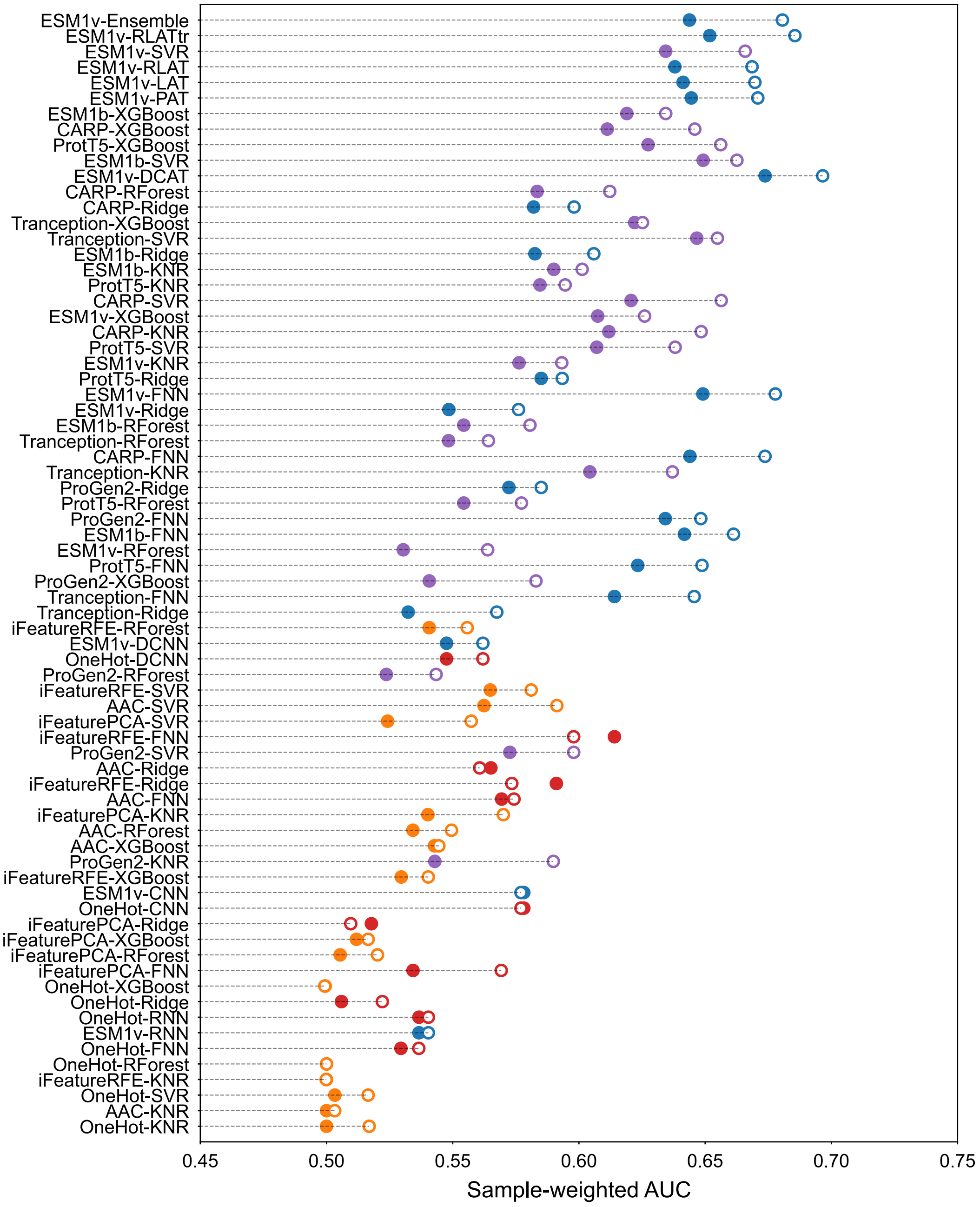


**Supplementary Figure 4.** Sample-weighted area under the ROC curve (AUC) of binned pHopt classes (acidic, neutral, alkaline) predicted by optimal models, evaluated on both the complete held-out pHopt testing set (n=1,971) and a subset of the testing set with less than 20% sequence identity to the training set (n=999).

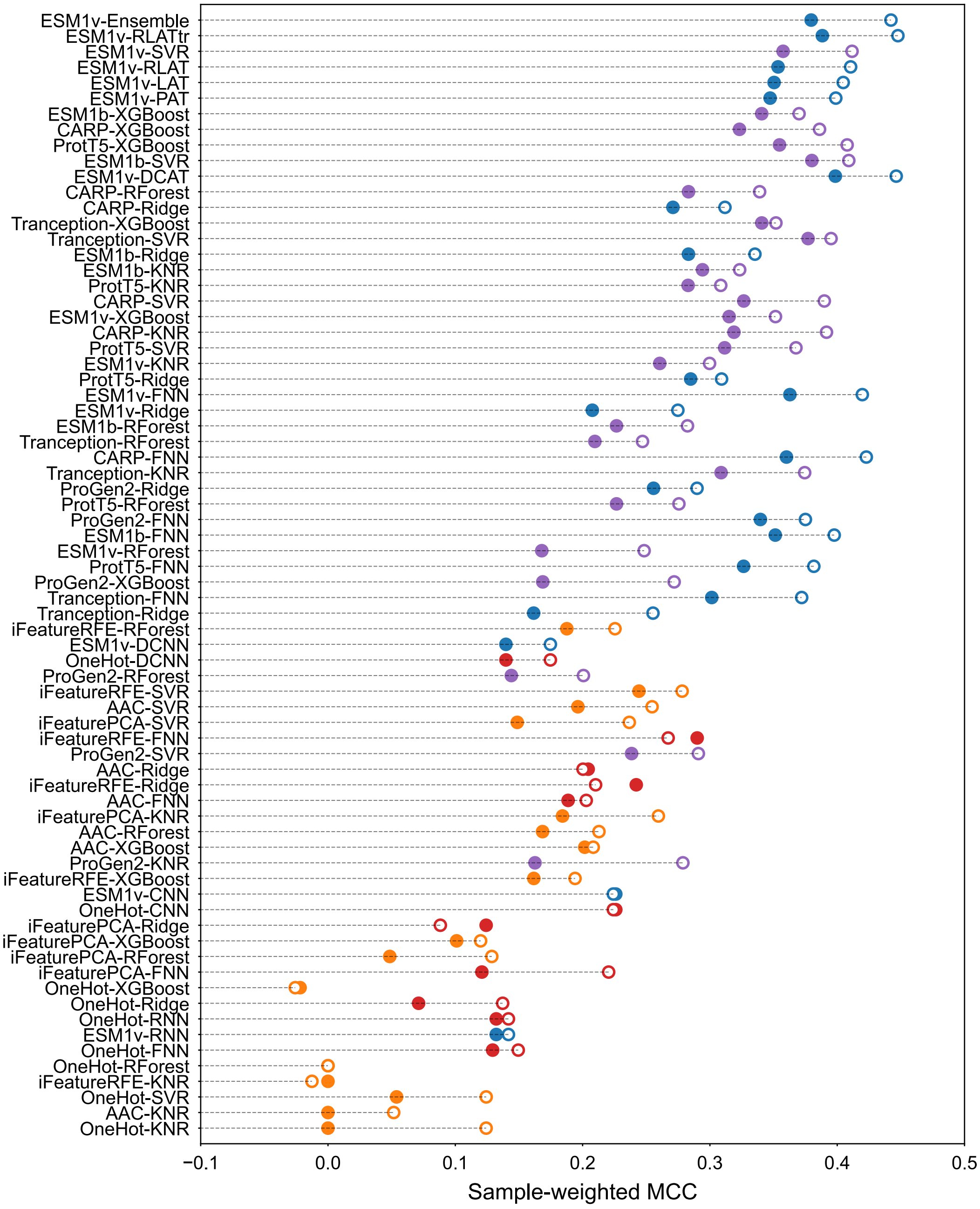


**Supplementary Figure 5.** Sample-weighted Matthew's correlation coefficient (MCC) of binned pHopt classes (acidic, neutral, alkaline) predicted by optimal models, evaluated on both the complete held-out pHopt testing set (n=1,971) and a subset of the testing set with less than 20% sequence identity to the training set (n=999).

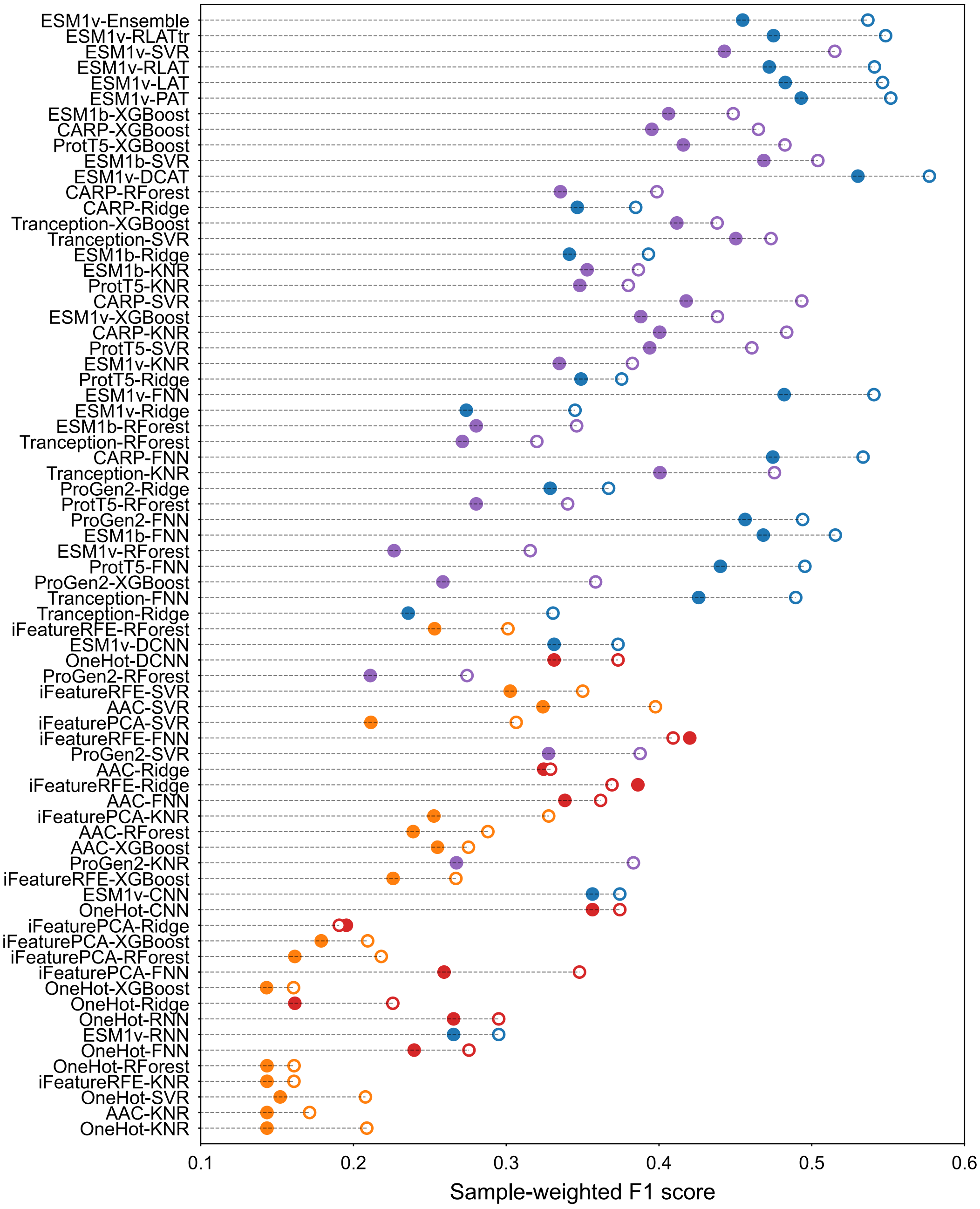


**Supplementary Figure 6.** Sample-weighted F1 score of binned pHopt classes (acidic, neutral, alkaline) predicted by optimal models, evaluated on both the complete held-out pHopt testing set (n=1,971) and a subset of the testing set with less than 20% sequence identity to the training set (n=999).

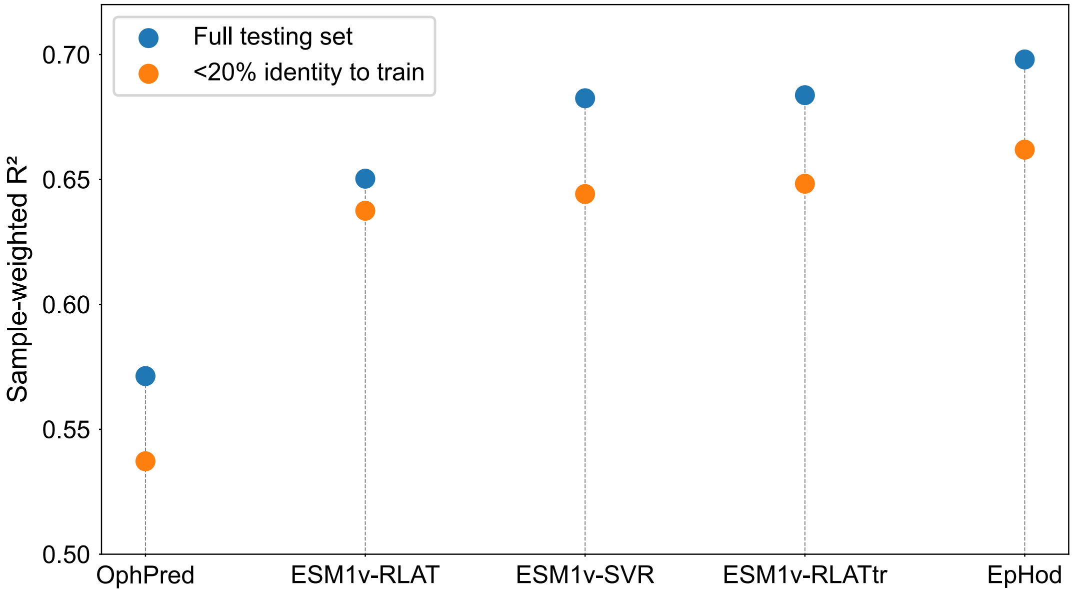


**Supplementary Figure 7**. Comparison of EpHod with OphPred.[1] EpHod is an ensemble of ESM1v-RLATtr and ESM1v-SVR.

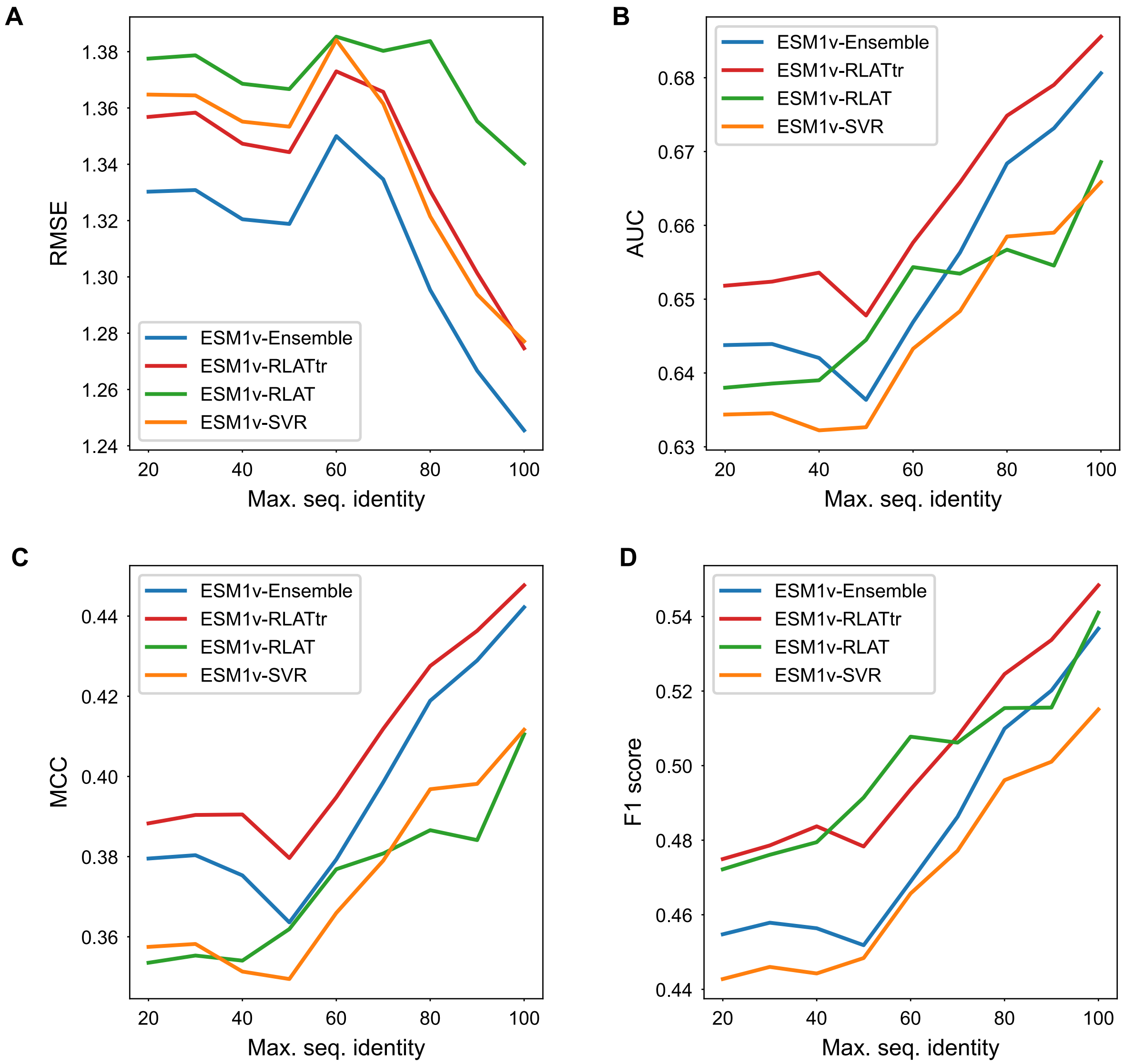


**Supplementary Figure 8**. Performance of the best-performing models on the testing set after applying maximum sequence identity thresholds. This evaluation was conducted on subsets of the testing set, where sequences exceeding the identity threshold were removed. For each point on the x-axes, sequences in the testing set with identity greater than the specific level compared with training sequences were removed, and the metric was evaluated on the remaining sequences.

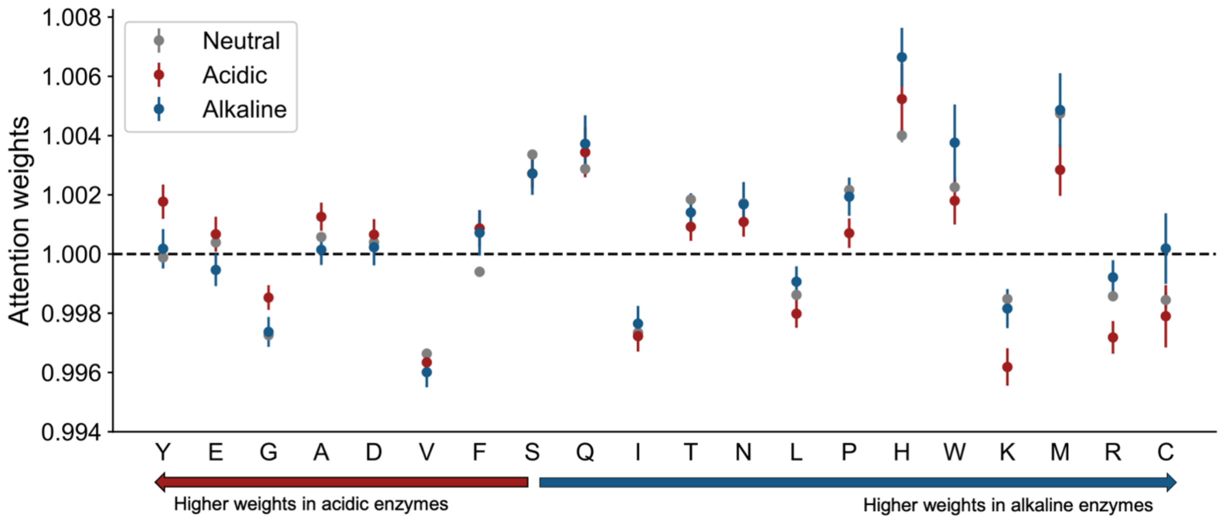


**Supplementary Figure 9**. Average attention weights of amino acid residues in the pHopt dataset (n=9,855). Weights were normalized to a mean value of 1.0 for each protein. Error bars indicate 95% confidence interval of the mean across the entire dataset. Overall, comparing acidic and alkaline enzymes, EpHod assigns increasingly greater weights to Ser, Phe, Val, Asp, Ala, Gly, Glu, and Tyr, in acidic enzymes (pHopt < 5), and increasingly greater weights to Gln, Ile, Thr, Asn, Leu, Pro, His, Trp, Lys, Met, Arg, and Cys in alkaline enzymes (pHopt > 9).

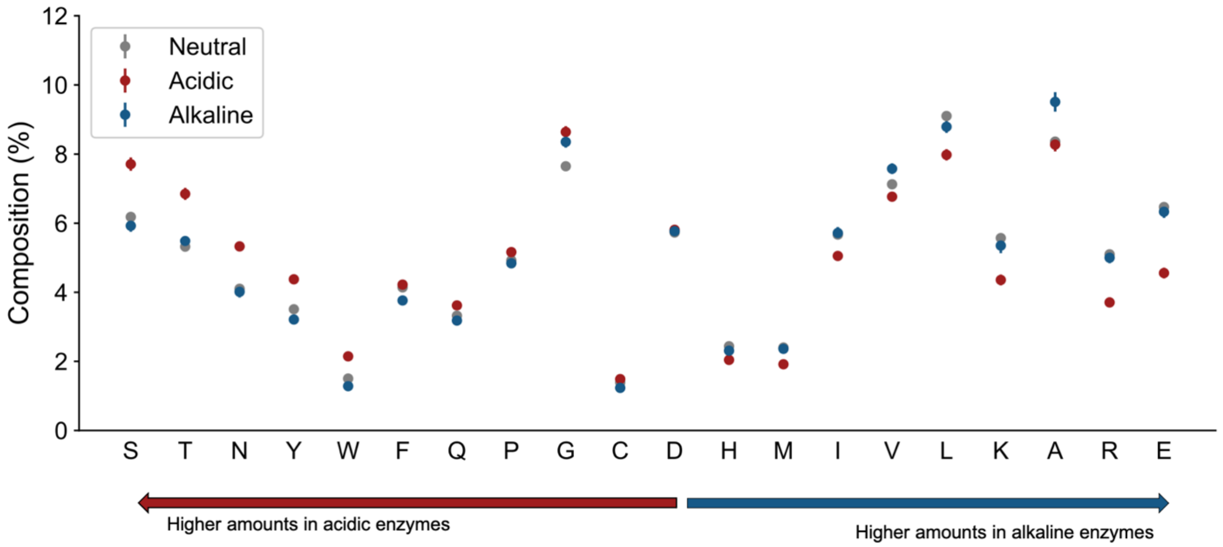


**Supplementary Figure 10**. Average composition of amino acids in the pHopt dataset (n=9,855). Error bars indicate 95% confidence interval of the mean across the entire dataset. Note that some error bars are too small to be seen. Overall, acidic enzymes have increasingly greater amounts of Asp, Cys, Gly, Pro, Gln, Phe, Trp, Tyr, Asn, Thr, and Ser, whereas alkaline enzymes have increasingly greater amounts of His, Met, Iso, Val, Leu, Lys, Ala, Arg, and Glu.

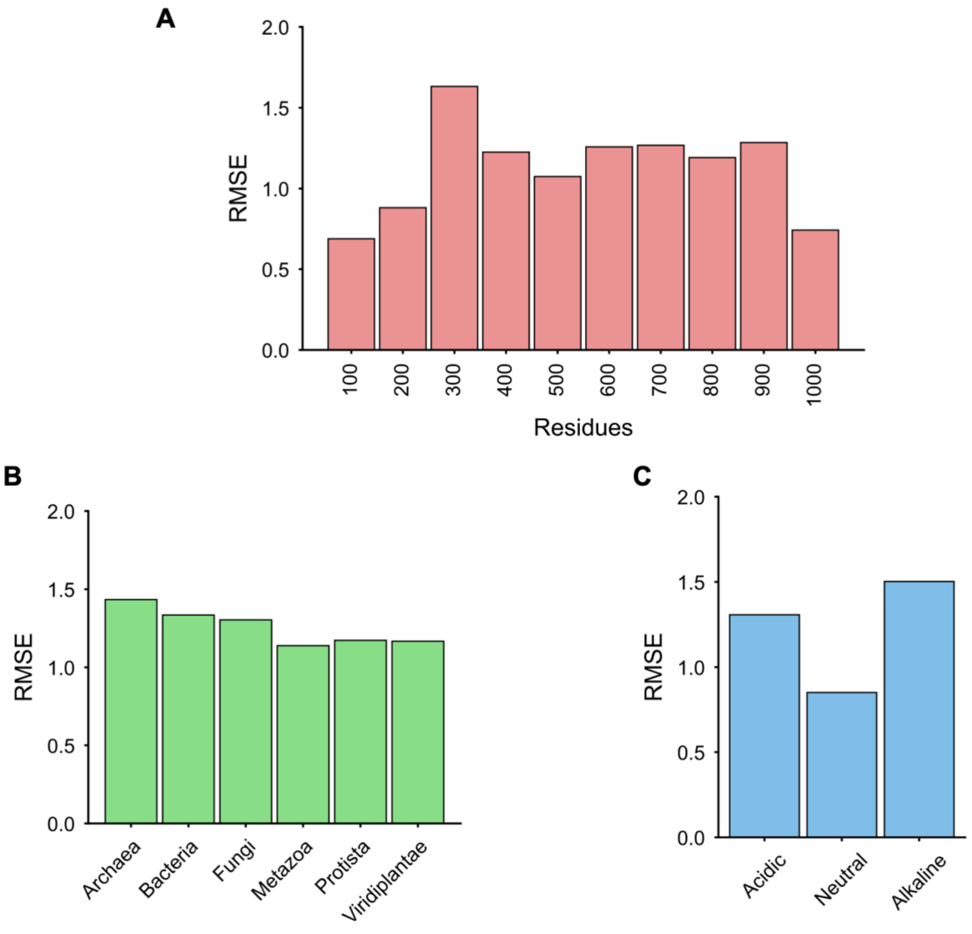


**Supplementary Figure 11.** Root mean squared error (RMSE) of ESM1v-RLAT $pH_{opt}$ predictions on testing set enzymes (n=1,971) split into subgroups according to **(A)** sequence length, **(B)** taxonomy, and **(C)** $pH_{opt}$ label class. RMSE values of less than 2.0 pH units are observed for all categories.

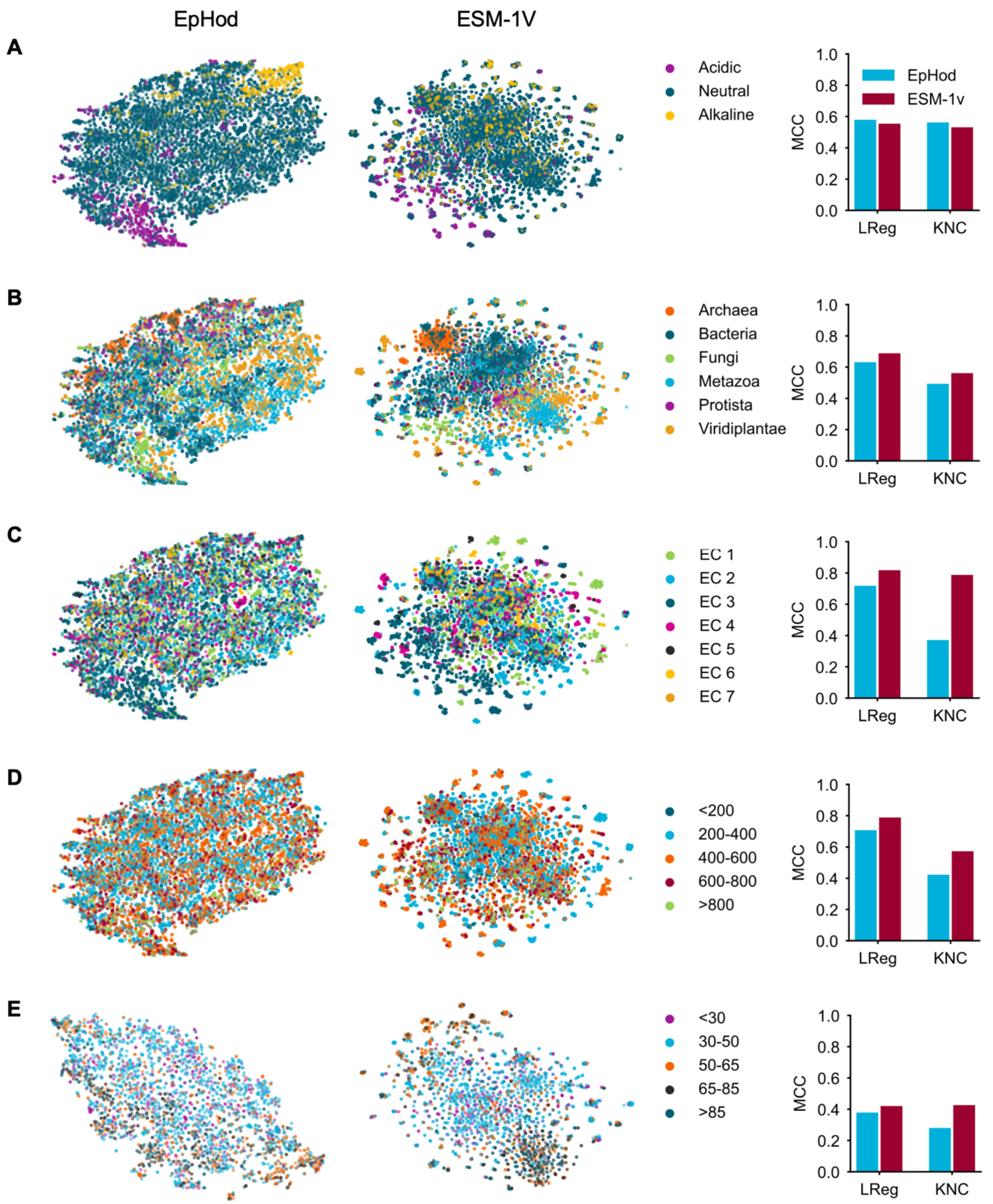


**Supplementary Figure 12**. Comparison of EpHod (ESM1v-RLATtr) and ESM-1v embeddings for supervised prediction tasks. EpHod embeddings are output of the final hidden dense layer (2,560-dim) and ESM-1v embeddings are the averaged output of the final transformer layer (1,280-dim). The figure presents a two-dimensional t-SNE visualization of the embeddings (left) and the Matthew's correlation coefficient (MCC) for multi-class classification using logistic regression (LReg) and K-neighbor classifiers (KNC)

with optimal hyperparameters based on the embeddings (right). Compared to ESM-1v, the marked restructuring of EpHod embeddings for improved pHopt prediction is apparent. Although ESM-1v generally outperforms EpHod on other tasks, EpHod embeddings still retain global informativeness for decent performance on the various tasks. (**A**) Optimal pH of 9,855 enzymes in the pHopt dataset (**B**) Optimal temperature ($T_{opt}$ in ºC) of 2,813 enzymes from Gado *et al.*[2] (**C**) Enzyme activity classes (EC numbers) of 9,855 enzymes in pHopt dataset (**D**) Organismal taxonomy of 9,855 proteins in pHopt dataset (**E**) Sequence length (number of residues) of 9,855 enzymes in pHopt dataset.

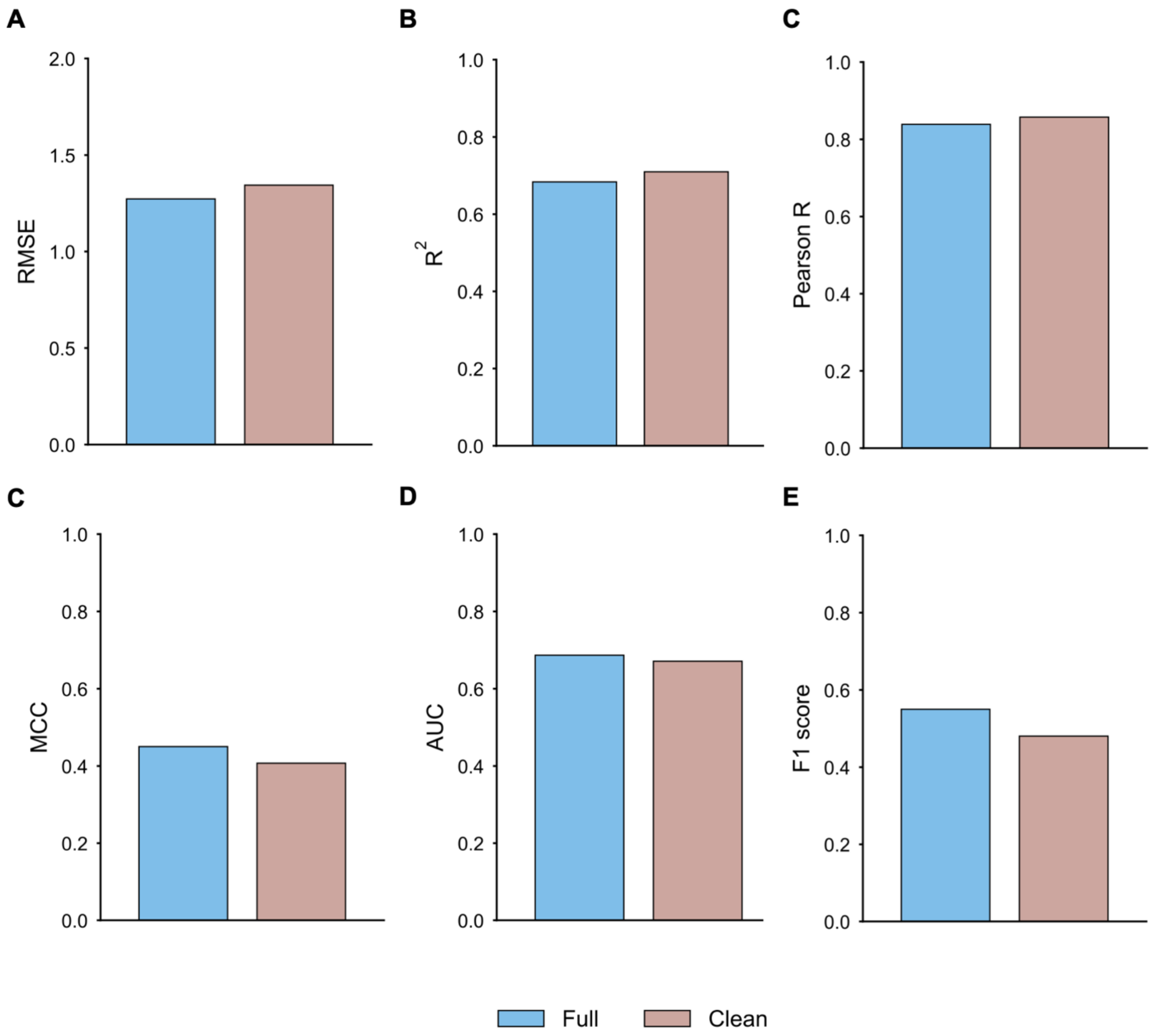


**Supplementary Figure 13.** Predictive performance of ESM1v-RLATtr on the full held-out testing set (n=1,971), and on a "clean" subset of the testing set (n=993) that contained only enzymes without the "assay at" annotation in BRENDA.[3] Entries with the "assay at" annotation were likely assayed at a single pH value, which may deviate significantly from the true pHopt and constitute noise in the dataset. However, EpHod demonstrates very similar performance on both full and "clean" datasets, suggesting that, in training, EpHod was robust to the experiment noise in the dataset arising due to the limited assay conditions, and learned general relationships between sequence and pHopt from the data.

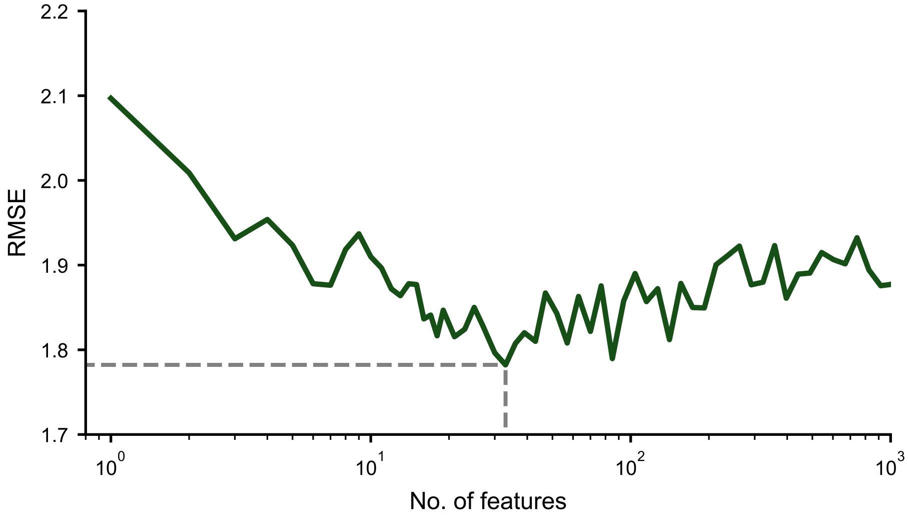


**Supplementary Figure 14.** Recursive feature elimination (RFE) approach to select an optimal subset of features from an initial pool of 5,494 features (iFeature).[4] The RFE process was implemented by discarding 10% of the features with the lowest random forest Gini importance scores in each iteration. The predictive performance (RMSE) was evaluated on the validation set, and the optimal number of features correspond to 33 features with RMSE of 1.78 (dashed lines).

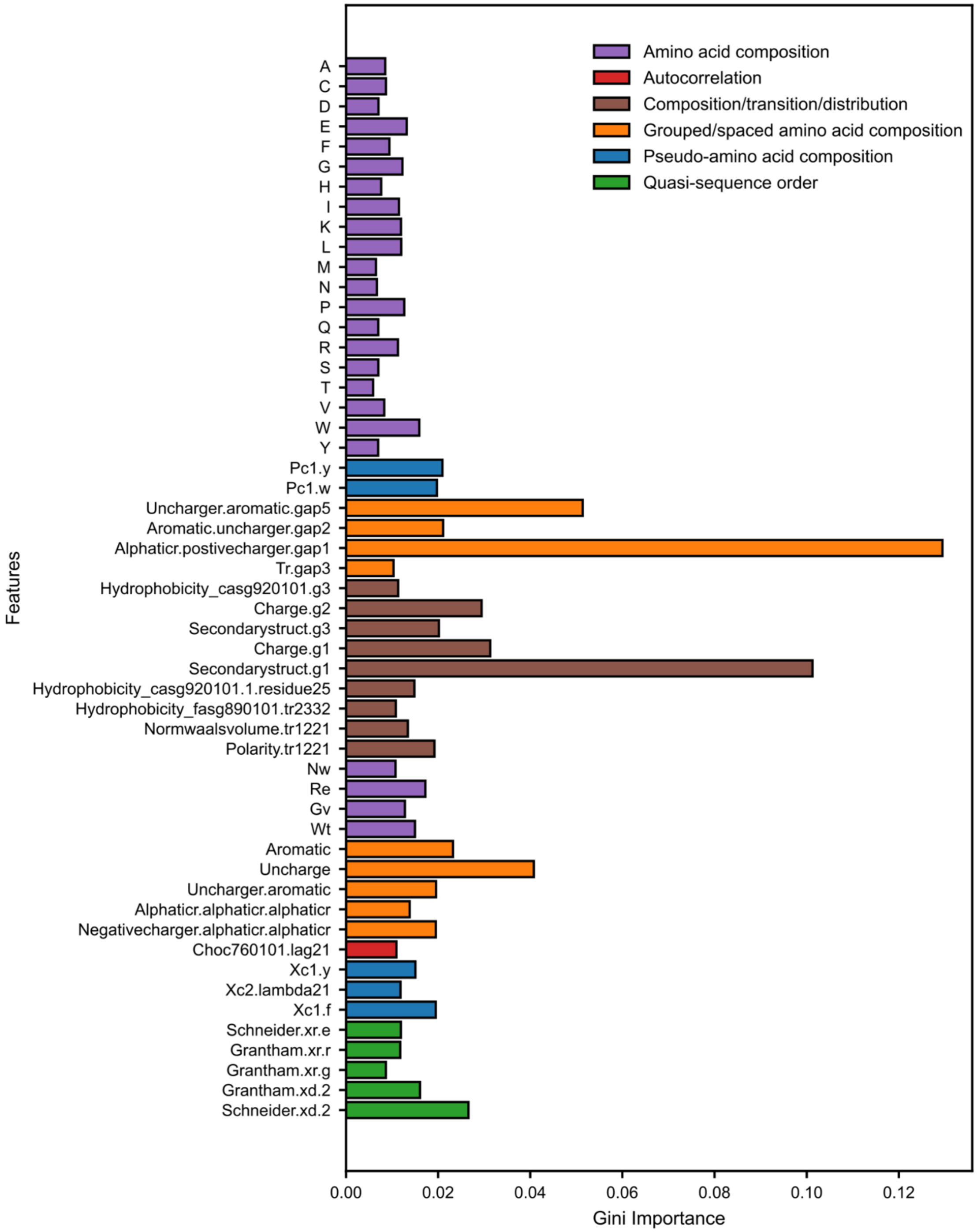


**Supplementary Figure 15**. Gini feature importances in trained random forest model for top 53 features selected from 5,494 iFeatures with recursive feature elimination (RFE).[4] The most important feature is *Aliphaticr.positivecharger.gap1*, which is the composition of the group defined by an aliphatic amino acid and a positively charged amino acid separated by a gap (any other amino acid).

**Supplementary Table 1**. Taxonomical distribution of enzymes in pHopt dataset (9,855 sequences).

| Kingdom | Frequency (%) |
|---|---|
| Archaea | 8.3 |
| Bacteria | 37.3 |
| Protista | 3.7 |
| Fungi | 10.6 |
| Viridiplantae | 16.4 |
| Metazoa | 23.6 |

**Supplementary Table 2**. Protein language models used to derive embeddings for predicting pHopt values. Embeddings were obtained from the final hidden layer of each model, and a supervised model was trained to predict pHopt values based on these embeddings.

| Name | Full model name | No. of Parameters | Embedding dimension | Training objective | Training sequences |
|---|---|---|---|---|---|
| ESM1v | ESM1v-1 (1 of 5 in ensemble) | 650M | 1,280 | Masked | Uniref90: 98M |
| ESM1b | ESM1b | 650M | 1,280 | Masked | Uniref50: 27M |
| CARP | CARP640 | 640M | 1,280 | Masked | Uniref50: 42M |
| ProtT5 | ProtT5-U50-XL | 3,000M | 1,024 | Masked | BFD100: 2,122M Uniref50: 45M |
| ProGen2 | ProGen2-base | 764M | 1,536 | Next token (autoregressive) | BFD30 Uniref90 |
| Tranception | Tranception-Large | 700M | 1,280 | Next token (autoregressive) | Uniref100: 250M |

**Supplementary Table 3**. Hyperparameter space of traditional machine learning models searched during model optimization. For each method, the hyperparameter space also includes five reweighting methods (See Figure 1B, main text). Hyperparameter names are consistent with the scikit-learn package.[5]

| **Method** | **Hyperparameter name** | **Search space** | **Description** |
|---|---|---|---|
| Ridge regression | alpha | 10^[-8, 8] | Strength of L2 regularization |
| K-neighbors regression (KNR) | n_neighbors | {1, 2, 5, 10, 20, 50, 100} | Number of nearest neighbors (k) |
| | weightby | {'uniform', 'distance'} | Method of neighbor weighting |
| Support vector regression (SVR) | kernel | {'poly', 'rbf'} | The kernel type used in the algorithm |
| | gamma | {'scale', 'auto'} | Method of computing kernel coefficient |
| | C | 10^[-5,5] | Inverse of regularization strength |
| Random forest (RForest) | n_estimators | {10, 20, 50, 100, 200, 500, 1000} | The number of trees |
| | criterion | {'mae', 'mse'} | Loss/error function |
| | max_features | {0.25, 0.5, 0.75, None} | Size of features to consider |
| | max_samples | {0.25, 0.5, 0.75, None} | Size of samples to consider |
| | max_depth | {5, 10, None} | Maximum depth of the tree |
| XGBoost | learning_rate | {0.01, 0.5, 0.1, 0.2, None} | Step size for boosting process |
| | reg_alpha | {0, 0.01, 0.1, 1, 10, 100, None} | L1 regularization weight |
| | reg_lambda | {0.01, 0.1, 1, 10, 100, None} | L2 regularization weight |
| | max_depth | {3, 5, 7, 9, 11, None} | Maximum depth of trees |
| | max_delta_step | {0, 1, 5, 10, None} | Maximum step for weight estimation |
| | min_child_weight | {1, 3, 5, 7, 10} | Minimum sum of instance weight |
| | n_estimators | {20, 50, 100, 200, 500, None} | Number of boosting rounds |

**Supplementary Table 4**. Hyperparameter space of neural network models searched during model optimization. For each method, the hyperparameter space also includes five reweighting methods (See Figure 1B, main text).

| **Method** | **Hyperparameter name** | **Search space** | **Description** |
|---|---|---|---|
| Feed-forward neural network (FNN) | hidden_dims | {128 256, 512, 1024} | Number of units in hidden layer |
| | num_layers | {1, 2, 3} | |
| | activation | {'relu', 'leaky_relu', 'elu', 'gelu'} | Activation function in hidden layer |
| | dropout | {0, 0.1, 0.25, 0.33, 0.5} | Hidden layers dropout fraction |
| | l2reg | {0, 1e-5, 1e-4, 1e-3, 1e-2} | L2 regularization (weight decay) |
| | learning_rate | {1e-4, 5e-4, 1e-3, 5e-3, 1e-2} | Initial learning rate |
| Convolution neural network (CNN) | num_conv_layers | {2,3,4,5,6} | Number of convolution layers |
| | kernel_size | {3, 5, 7} | Kernel size of 1D convolution |
| | conv_dropout | {0, 0.1, 0.25, 0.33, 0.5} | Dropout in convolution layers |
| | pool_type | {'max', 'average'} | Max or average pooling |
| | dense_dim | {32, 64, 128, 256, 512, 1024} | Number of units in dense layers after convolution layers |
| | num_dense_layers | {1, 2, 3, 4} | Number of dense layers of same unit size. |
| | dense_dropout | {0, 0.1, 0.25, 0.33, 0.5} | Dropout in dense hidden layers |
| | activation | {'relu', 'leaky_relu', 'elu', 'gelu'} | Activation function in hidden layers |
| | learning_rate | {1e-5, 5e-5, 1e-4, 5e-4, 1e-3, 5e-3} | Initial learning rate in optimization |
| | l2reg | {0, 1e-5, 1e-4, 1e-3, 1e-2} | L2 regularization (weight decay) |
| Dlated convolution neural network (DCNN) | conv_channel | {24, 32, 48, 64, 96} | Number of channels |
| | num_blocks | {1, 2, 3, 4, 6} | Number of dilated CNN blocks |
| | block_dropout | {0, 0.1, 0.25, 0.33, 0.5} | Dropout after convolution block |
| | attention_dim | {32, 64, 128, 256, 512, 1024} | Dimension of convolution layer to learn attention weights |
| | attention_kernel_size | {3, 5, 7, 9, 11, 13} | Kernel size of attention convolution |
| | dense_dim | {32, 64, 128, 256, 512, 1024} | Number of units in hidden dense layer |
| | dense_dropout | {0, 0.1, 0.25, 0.33, 0.5} | Dropout in dense hidden layers |
| | activation | {'relu', 'leaky_relu', 'elu', 'gelu'} | Activation function in hidden layers |
| | learning_rate | {1e-5, 5e-5, 1e-4, 5e-4, 1e-3, 5e-3} | Initial learning rate |
| | l2reg | {0, 1e-5, 1e-4, 1e-3, 1e-2} | L2 regularization (weight decay) |
| Recurrent neural network GRU (RNN) | gru_dim | {64, 128, 256, 512, 1024} | Feature dimension of GRU |
| | conv_downsample | {1, 2, 4, 8} | Stride size of convolution layer to downsample input |
| | conv_dropout | {0, 0.1, 0.25, 0.33, 0.5} | Dropout after convolution layer |
| | dense_dim | {64, 128, 256, 512, 1024} | Number of units in hidden dense layer |
| | dense_dropout | {0, 0.1, 0.25, 0.33, 0.5} | Dropout in dense hidden layers |
| | activation | {'relu', 'leaky_relu', 'elu', 'gelu'} | Activation function in hidden layers |
| | learning_rate | {1e-5, 5e-5, 1e-4, 5e-4, 1e-3, 5e-3} | Initial learning rate |

| | | | |
|---|---|---|---|
| | l2reg | {0, 1e-5, 1e-4, 1e-3, 1e-2} | L2 regularization (weight decay) |
| Light attention (LAT)<br><br>or<br><br>Perceptive light attention (PAT)<br><br>or<br><br>Dilated convolution with light attention (DCAT)<br><br>or<br><br>Residual light attention (RLAT) | kernel_size | {3, 5, 7, 9, 11, 13} | Kernel size of attention convolution layer |
| | conv_dropout | {0, 0.1, 0.25, 0.33, 0.5} | Dropout after convolution/attention layer. Not used in RLAT. |
| | perceptive | {True, False} | If True, concatenate embeddings with a repeated average-pooled tensor. Only used in PAT. |
| | dense_dim | {32, 64, 128, 256, 512, 1024} | Number of units in hidden dense layer. |
| | dense_dropout | {0, 0.1, 0.25, 0.33, 0.5} | Dropout after dense layers. |
| | conv_dim | {128, 256, 512} | Number of channels in convolution layers after the attention layers. Only used in DCAT. |
| | activation | {'relu', 'leaky_relu', 'elu', 'gelu'} | Activation function in hidden layers |
| | learning_rate | {5e-6, 1e-5, 5e-5, 1e-4, 5e-4, 1e-3, 5e-3} | Initial learning rate in optimization |
| | l2reg | {0, 1e-5, 1e-4, 1e-3, 1e-2} | L2 regularization (weight decay) |

**Supplementary Table 5.** Number of models trained with per-protein vector representations during hyperparameter optimization. Language model embeddings (*italicized*) were averaged and one-hot representations were flattened to derive a vector for each protein. A total of 9,550 model instances were trained.

| | **Ridge** | **KNR** | **SVR** | **RForest** | **XGBoost** | **FNN** |
|---|---|---|---|---|---|---|
| AAC | 85 | 70 | 200 | 200 | 200 | 200 |
| iFeature (PCA) | 85 | 70 | 200 | 200 | 200 | 200 |
| iFeature (RFE) | 85 | 70 | 200 | 200 | 200 | 200 |
| One Hot | 85 | 70 | 200 | 200 | 200 | 200 |
| *ESM-1v* | 85 | 70 | 200 | 200 | 200 | 200 |
| *ESM-1b* | 85 | 70 | 200 | 200 | 200 | 200 |
| *ProtT5* | 85 | 70 | 200 | 200 | 200 | 200 |
| *ProGen2* | 85 | 70 | 200 | 200 | 200 | 200 |
| *Tranception* | 85 | 70 | 200 | 200 | 200 | 200 |
| *CARP* | 85 | 70 | 200 | 200 | 200 | 200 |

**Supplementary Table 6.** Number of neural network models trained with per-residue representations during hyperparameter optimization. A total of 2,000 model instances were trained.

| | **CNN** | **DCNN** | **RNN** | **LAT** | **RLAT** | **PAT** | **DCAT** |
|---|---|---|---|---|---|---|---|
| One Hot | 200 | 200 | 200 | N/A | N/A | N/A | N/A |
| *ESM-1v* | 200 | 200 | 200 | 200 | 200 | 200 | 200 |

**Supplementary Table 7**. Description of 5,494 features generated by the iFeature package used in predicting pHopt in this work.[4]

| **Feature** | **Description** | **Count** |
|---|---|---|
| AAC | Amino acid composition | 20 |
| CKSAAP | Composition of k-spaced amino acid pairs | 2,400 |
| DPC | Dipeptide composition | 400 |
| DDE | Dipeptide deviation from expected mean | 400 |
| GAAC | Grouped amino acid composition | 5 |
| CKSAAGP | Composition of k-spaced amino acid group pairs | 150 |
| GDPC | Grouped dipeptide composition | 25 |
| GTPC | Grouped tripeptide composition | 125 |
| Moran | Moran autocorrelation | 240 |
| Geary | Geary autocorrelation | 240 |
| NMBroto | Normalized Moreau-Broto autocorrelation | 240 |
| CTDC | Composition | 39 |
| CTDT | Transition | 39 |
| CTDD | Distribution | 195 |
| CTriad | Conjoint triad | 343 |
| KSCTriad | Conjoint k-spaced triad | 343 |
| SOCNumber | Sequence-order-coupling number | 60 |
| QSOrder | Quasi-sequence-order descriptors | 100 |
| PAAC | Pseudo-amino acid composition | 50 |
| APAAC | Amphiphilic pseudo-amino acid composition | 80 |
| Total | | 5,494 |

#### Supplementary References


1. Zaretckii, M., Buslaev, P., Kozlovskii, I., Morozov, A. & Popov, P. Approaching Optimal pH Enzyme Prediction with Large Language Models. *ACS Synth. Biol.* (2024) doi:10.1021/acssynbio.4c00465.
2. Gado, J. E., Beckham, G. T. & Payne, C. M. Improving enzyme optimum temperature prediction with resampling strategies and ensemble learning. *J. Chem. Inf. Model.* **60**, 4098–4107 (2020).
3. Li, G. *et al.* Performance of Regression Models as a Function of Experiment Noise. *Bioinform Biol Insights* **15**, 11779322211020315 (2021).
4. Chen, Z. *et al.* iFeature: a Python package and web server for features extraction and selection from protein and peptide sequences. *Bioinformatics* **34**, 2499–2502 (2018).
5. Pedregosa, F. *et al.* Scikit-learn: machine learning in Python. *Journal of Machine Learning Research* **12**, 2825–2830 (2011).
